# Embryonic expression and developmental RNAi phenotypes of genes associated with juvenile hormone and ecdysone pathways in *Tribolium castaneum*

**DOI:** 10.64898/2026.11.15.766745

**Authors:** Alessandra da Silva de Alvarenga, Lupis Ribeiro Gomes Neto, Bruno Rodrigues, Daniele Santos, Rodrigo Nunes-da-Fonseca

## Abstract

Juvenile hormone (JH) and ecdysteroids coordinate insect molting and metamorphosis, but the embryonic deployment of genes associated with these pathways remains incompletely characterized. We examined the JH biosynthetic genes *Tc-jhamt* and *Tc-epox* (*CYP15A1*), the ecdysone receptor gene *Tc-EcR* and the micropeptide gene *mille-pattes* (*Tc-mlpt*) in the red flour beetle *Tribolium castaneum*. Whole-mount in situ hybridization revealed a discrete *Tc-epox* domain in the differentiated blastoderm followed by bilateral domains in the early germ rudiment, paired *Tc-jhamt* domains in three consecutive trunk segments of the extended germ band, patterned *Tc-mlpt* expression and weak, broad *Tc-EcR* signal. After parental RNAi, mean hatching was 9.0% for *Tc-mlpt* and 26.0% for *Tc-EcR*, compared with 74.0% in water controls, whereas hatching after *Tc-epox* and *Tc-jhamt* RNAi (74.5% and 73.0%) was close to control values. Offspring of *Tc-epox*- and *Tc-jhamt*-injected females nevertheless included embryos with anterior or germ-band defects and unhatched larvae with malformed cuticles, and unhatched *Tc-EcR* offspring showed sparse or fragmented cuticle. After larval RNAi, mean mortality by day 18 was 98.8% for *Tc-mlpt* and 91.3% for *Tc-EcR* (controls, 8.8%), and affected larvae failed to shed the old cuticle. These results extend the spatial map of JH biosynthetic gene expression to early stages of a holometabolous embryo and identify a role for *Tc-mlpt* in larval ecdysis in beetles.

## Introduction

The evolution of complete metamorphosis was one of the key innovations in insect history. Holometabolous insects, which pass through larval, pupal and adult stages, comprise most described insect species, whereas hemimetabolous insects hatch as nymphs that gradually acquire adult features, and the wingless ametabolous orders (Archaeognatha and Zygentoma) develop without metamorphosis [1–3]. How the holometabolous larva and pupa arose from a hemimetabolous ancestor has long been debated. The classical view holds that the larva corresponds to the nymphal instars and the pupa to a modified final nymphal stage, whereas the pronymph hypothesis proposes that the larva derives from the pronymph, a transient embryonic stage of hemimetabolous insects, and that the nymphal instars were compressed into the pupa [1, 4–6]. Both views place changes in the timing of hormonal signals, and particularly of embryonic endocrine events, at the center of this transition [2, 7–9].

Two hormones govern insect molting and metamorphosis. Ecdysteroids, acting through the ecdysone receptor (EcR), trigger each molt, whereas juvenile hormone (JH), acting through its receptor Methoprene-tolerant (Met), determines the developmental identity of the molt [10]. Downstream of these receptors, the MEKRE93 pathway connects JH and ecdysone signaling to stage specification: JH induces *Krüppel homolog 1* (*Kr-h1*), which represses the adult specifier *E93*, while *broad-complex* (*br-C*) specifies the pupal stage [11–15]. The anti-metamorphic roles of Met and Kr-h1 are conserved from cockroaches to beetles [16, 17], but their requirement is stage-dependent: in the bug *Pyrrhocoris apterus* and in JH-deficient silkworms, the first larval instars develop normally without JH signaling, which becomes essential only once larvae acquire competence to metamorphose [18–20].

Recent work in ametabolous and hemimetabolous species has shifted attention to the embryo as the likely ancestral site of JH action. In the firebrat *Thermobia domestica*, JH titers peak in late embryogenesis, and loss- and gain-of-function experiments show that JH promotes the transition from morphogenesis to terminal differentiation of embryonic tissues [21]. Knockout of *JHAMT*, *CYP15A1*, *Met* or *Kr-h1* in *Thermobia* causes embryonic lethality, with defects in post-dorsal-closure development, leg maturation and hatching, and similar effects were found in the cricket *Gryllus bimaculatus* [22]. In hemimetabolous embryos, JH appears with the formation of the first nymphal cuticle, premature JH induces precocious nymphal features [23], and JH signaling is required for normal embryogenesis of *Blattella germanica* [24]. Embryonic expression of *br-C* in *Thermobia* and in the milkweed bug *Oncopeltus fasciatus*, absent in holometabolous embryos [25], and the reduction of embryonic *E93* expression [26] have been proposed as molecular changes underlying the origin of the holometabolous larva. These findings suggest that the embryonic functions of JH preceded its post-embryonic anti-metamorphic role [21, 22, 27]. In holometabolous insects, however, the embryonic roles of JH appear more variable: silkworm embryos lacking JH synthesis or perception complete morphogenesis [19], whereas parental RNAi against *Met* blocks embryonic development in *Tribolium castaneum* [28].

The final steps of JH biosynthesis are the methylation and epoxidation of sesquiterpenoid precursors by JH acid O-methyltransferase (JHAMT) and by CYP15 P450 enzymes [29, 30]. Their order and the enzymes involved vary among insects: CYP15A1 epoxidizes methyl farnesoate in cockroaches and locusts [30, 31], lepidopteran CYP15C1 epoxidizes farnesoic acid and is required for JH synthesis in *Bombyx mori* [18], and higher dipterans lack *CYP15* and rely on Cyp6g2 [32]. JHAMT genes have undergone lineage-specific duplications [33], and in *Drosophila* loss of *jhamt* is compensated by methyl farnesoate, which itself acts through the JH receptors [34]. The sesquiterpenoid pathway predates insects: in the crustacean *Daphnia magna*, *Jhamt* is transiently activated around hatching and controls the duration of embryogenesis [35], and epoxidation of JH has been proposed as an insect innovation [36]. In *T. castaneum*, Tc-MT3 was characterized as JHAMT, whereas the related Tc-MT1 and Tc-MT2 did not methylate the tested substrates [37], and Tc-CYP15A1 epoxidizes both farnesoic acid and methyl farnesoate [38]. Transcripts of both enzymes are detected in *Tribolium* embryos [37, 38], but their spatial distribution, which could indicate sites of JH synthesis before the corpora allata differentiate, has not been described.

Ecdysteroid signaling also operates before hatching. In *Drosophila melanogaster*, it is required for mid-embryonic morphogenesis and organ development [39, 40], and ecdysteroid synthesis is required for gastrulation in *Daphnia* embryos [35]. In *Blattella*, EcR-A is required for nymphal molting [41], and in *Tribolium*, EcR isoforms play distinct roles in molting and metamorphosis [42]. In *Tribolium* embryos reared at 25°C, ecdysteroid titers show a weak peak at about 40 h, when the germ band is extended, and a strong peak at about 120 h, at dorsal closure, before hatching at about 153 h; the ecdysone-inactivating enzyme Cyp18a1 is required for dorsal closure, and a regulatory allele of *Cyp18a1* that advances the late peak shortens development [43, 44].

One link between ecdysone signaling and morphogenesis is provided by the polycistronic micropeptide gene *mille-pattes*/*polished rice*/*tarsal-less* (*mlpt*/*pri*/*tal*), the best-characterized example of the small open reading frame (sORF) genes, which are now recognized as a widespread source of regulatory peptides in insects [45, 46]. In *Drosophila*, *pri*/*tal* encodes 11–32-amino-acid peptides that control epidermal differentiation and leg development [47, 48] by inducing the proteasome-dependent processing of the transcription factor Shavenbaby (Svb) into an activator [49, 50]. Pri peptides mediate ecdysone-dependent developmental timing and embryonic organogenesis [51–53], act pleiotropically during leg morphogenesis [54] and function as a cell-intrinsic timer of fate induction [55]. Their functions extend beyond embryogenesis: *tal* is required for tarsal segmentation of the adult leg [47, 56], the ecdysone/Pri axis represses pupal cuticle genes throughout the larval stages to allow formation of a soft larval cuticle [57], and *pri* acts downstream of the ecdysone-response gene *ftz-f1* in the adult male accessory gland [58]. In *Tribolium*, *mlpt* behaves as a gap-like segmentation gene and controls appendage identity through the Mlpt/Ubr3/Svb module, whose embryonic patterning role is conserved in the wasp *Nasonia vitripennis* and in hemipterans [59, 60]. In the kissing bug *Rhodnius prolixus*, the *mlpt* mRNA encodes, in addition to the conserved LDPTG-containing peptides, a peptide of approximately 80 amino acids (smHemiptera) that is conserved among hemipterans but absent from other insect orders [45, 61]. *Rhodnius mlpt* regulates thoracic versus abdominal segmentation, leg development and head formation in the embryo [61] and is essential for nymphal molting and digestive physiology, with *mlpt* knockdown increasing the expression of *JHAMT*-like, *EPOX*-like and *Kr-h1* transcripts [62]. In the mosquito *Aedes aegypti*, *mlpt* transcripts are maternally provided [63]. In *Tribolium*, RNAi against *mlpt* in the last larval instar causes lethality just before pupation without affecting wing-disc proliferation [64], but whether *mlpt* is required for the larval–larval molts has not been examined.

*T. castaneum* combines a holometabolous life cycle with short-germ embryogenesis [65, 66] and a robust systemic RNAi response at both parental and larval stages [67, 68], making it well suited to examine how JH and ecdysone pathway genes are deployed in a holometabolous embryo and to compare them with ametabolous and hemimetabolous species. Here, we describe the spatial and temporal expression of *Tc-jhamt*, *Tc-epox*, *Tc-EcR* and *Tc-mlpt* during embryogenesis, place the biosynthetic enzymes in a phylogenetic context and analyze the embryonic, cuticular and larval phenotypes produced by parental and larval RNAi against these genes.

## Results

### Phylogenetic position of the target genes and specificity of probes

To place the target genes in an arthropod context, we searched 25 RefSeq arthropod proteomes and identified candidate homologs by reciprocal best hits (Fig. S1; Table S1). In the JHAMT tree (43 sequences, 176 aligned positions, LG+I+G4), Tc-JHAMT (MT3), Tc-MT1 and Tc-MT2 grouped with a single *Tenebrio molitor* JHAMT-like protein (SH-aLRT/UFBoot 94.2/97), and Tc-MT1 and Tc-MT2 were more closely related to each other and to the *Tenebrio* protein than to Tc-JHAMT (99.4/100) (Fig. S1a). To date these duplications, we repeated the analysis with 23 beetle genomes, including four other tenebrionids (*Tribolium madens*, *Gnatocerus cornutus*, *Zophobas morio* and *T. molitor*), and six other insects (124 sequences, 204 positions, Q.yeast+I+R4; Fig. S1d). Tenebrionid JHAMT-like proteins formed a single clade (98.1/100) that split into two lineages. One contained Tc-JHAMT and single orthologs from *T. madens*, *G. cornutus*, *Z. morio* and *T. molitor* (93.4/95); the other contained Tc-MT1, Tc-MT2 and paralogs from *T. molitor* and *Z. morio* (97.8/98). The duplication that separated Tc-JHAMT from the MT1/MT2 lineage therefore preceded the divergence of *Tribolium*, *Tenebrio* and *Zophobas*. Tc-MT1 and Tc-MT2 each had a one-to-one ortholog in *T. madens* (100/100), whereas their closest *Tenebrio* relative was a single protein (72.8/64), which places the MT1–MT2 duplication in the *Tribolium* lineage, after its separation from *Tenebrio* and before the divergence of *T. castaneum* and *T. madens*. The MT1/MT2 lineage expanded independently in *Z. morio* (17 paralogs) and *T. molitor* (6 paralogs), and no member of it was found in the *G. cornutus* annotation. Only Tc-MT3 is known to methylate JH precursors [37]. Leaf beetles (*Leptinotarsa*, *Diabrotica*) carried independent JHAMT duplicates, whereas dipteran, lepidopteran and hymenopteran JHAMTs formed separate, well-supported clades. In the CYP15 tree (31 sequences, 369 positions, LG+I+G4), Tc-CYP15A1 grouped with the coleopteran CYP15 sequences (93.6/98), and CYP15-related sequences, in some species more than one, were recovered from most insect orders and from crustaceans; sequence similarity alone does not establish epoxidase activity for these proteins (Fig. S1b). Both *Tribolium* EcR isoforms grouped within a coleopteran EcR clade (Fig. S1c).

Because JHAMT paralogs are closely related, we screened the probe and dsRNA templates against all 26,628 RefSeq transcripts of *T. castaneum* (Fig. S2; Table S2). The *Tc-mlpt* (477 bp), *Tc-epox* (764 bp) and *Tc-jhamt* (559 bp) templates had no exact 21-nt match to transcripts of other genes, and their best alignments to non-target transcripts did not exceed 66% identity; the *Tc-jhamt* template was 64% identical to its closest paralog, *Tc-mt1*. The *Tc-EcR* template (631 bp) spans the 5′ region specific to isoform A and part of the region shared by all isoforms, and is therefore expected to target both EcR-A and EcR-B. Within its DNA-binding domain, a 26–28-nt stretch is identical to transcripts of *E75* (two annotated loci) and *HR38*. Conventional RT-PCR amplified products of the expected sizes for all targets from embryonic cDNA (Fig. S3).

### Distinct spatial expression during embryogenesis

Whole-mount *in situ* hybridization revealed distinct patterns for the four genes (Fig. 1). *Tc-mlpt* was expressed in transverse domains from the blastoderm stage onwards, followed by anterior and posterior domains during germ-band formation and segmental expression in later embryos (Fig. 1a–d), in agreement with previous descriptions [59, 60]. *Tc-EcR* produced a weak, broadly distributed signal in blastoderm and germ-band embryos (Fig. 1e–g).

**Fig. 1.**
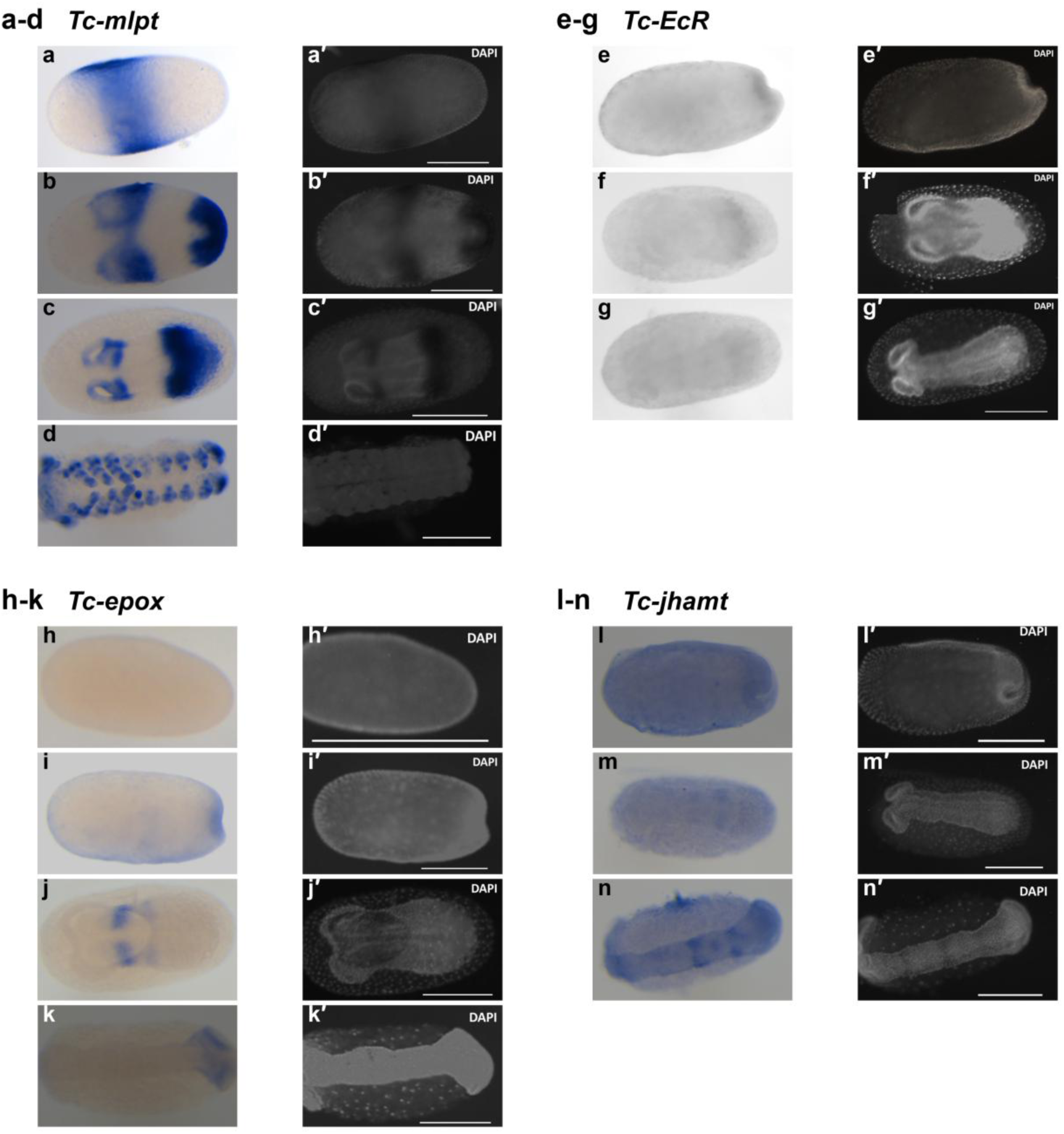
E**m**bryonic **expression of *Tc-mlpt*, *Tc-EcR*, *Tc-epox* and *Tc-jhamt*.** Whole-mount *in situ* hybridization (NBT/BCIP; left) and corresponding DAPI staining (right; primed labels). (a–d) Tc-mlpt is expressed in transverse domains in the blastoderm (a), in anterior and posterior domains during germ-band formation (b, c) and in segmental domains, including the appendage primordia, of the extended germ band (d). (e–g) Tc-EcR shows weak, broad signal in blastoderm (e) and germ-band embryos (f, g); colour images were not available for this probe. (h–k) Tc-epox: no detectable signal in the early blastoderm (h), signal at the posterior pit (i), paired bilateral domains in the forming germ rudiment (j) and a bilateral domain at the posterior region of an extending germ band (k). (l–n) Tc-jhamt: weak, non-localized signal in the blastoderm (l) and early germ band (m), and transverse, segmentally reiterated domains in the trunk of the extended germ band (n). Scale bars, 200 µm

*Tc-epox* and *Tc-jhamt* were analyzed in individually imaged embryos and in fields containing embryos at several stages, which were staged by DAPI morphology (*Tc-epox*, n = 17 embryos; *Tc-jhamt*, n = 20). *Tc-epox* signal was not detected in the early blastoderm (Fig. 1h). In the differentiated blastoderm, a small, sharply delimited domain appeared at approximately 60% egg length (Fig. S4a; visible in 3 of 5 embryos at the blastoderm and posterior-pit stages), and signal was present in the invaginating posterior region during formation of the posterior pit (Fig. 1i; Fig. S4b). As the germ rudiment formed, *Tc-epox* was expressed in a pair of bilateral domains (Fig. 1j; Fig. S4c), a pattern observed in all embryos at this stage. During germ band elongation, a bilateral domain at the posterior region of the germ band was observed (Fig. 1k; Fig. S4d).

*Tc-jhamt* signal was weak and not localized in blastoderm embryos (Fig. 1l; Fig. S4e; n = 5). In early germ bands, it was weak and diffuse, with enrichment in the posterior growth zone in some embryos (Fig. 1m; Fig. S4f; n = 4). During germ-band elongation, a single transverse stripe appeared in the trunk (Fig. S4g), and in extended germ bands *Tc-jhamt* was expressed in pairs of bilateral, lateral domains in three consecutive segments (Fig. 1n; Fig. S4h); stripes or paired domains were present in 9 of 11 embryos at these stages. The domains were spaced 55–60 µm apart, matching the width of one segment measured from the segmental furrows in the corresponding DAPI images (55–65 µm), and were located in the middle of the trunk, at a similar position to the single stripe of younger embryos. Their segmental identity remains to be determined with segment-specific markers.

### Temporal expression of *Tc-mlpt*, *Tc-epox* and *Tc-jhamt*

RT-qPCR across five collection intervals (0–6, 6–12, 12–24, 24–48 and 48–72 h after egg laying) revealed contrasting profiles (Fig. 2). Geometric means of *Tc-mlpt* rose from 1.10 at 0–6 h to 44.3 at 12–24 h, returned to basal levels at 24–48 h (1.08) and increased moderately at 48–72 h (6.6). *Tc-epox* increased throughout embryogenesis, from 2.0 at 0–6 h to 31.2 at 24–48 h and 252 at 48–72 h. *Tc-jhamt* remained between 1.0 and 2.1 during the first four intervals and reached 74.8 at 48–72 h, a 35-fold increase relative to 24–48 h. The late rise of both biosynthetic genes is thus temporally separated from the early *Tc-mlpt* peak, and their localized domains (Fig. 1) were already detectable at stages when whole-embryo transcript levels were low.

**Fig. 2.**
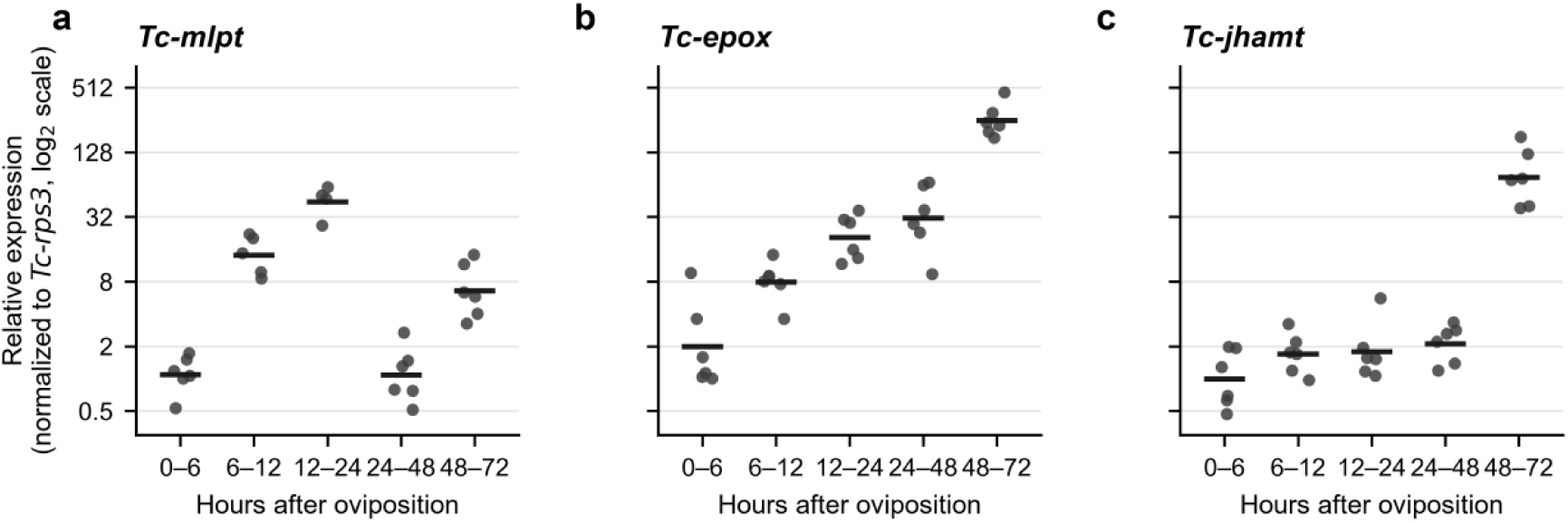
T**e**mporal **expression of *Tc-mlpt*, *Tc-epox* and *Tc-jhamt* during embryogenesis.** RT-qPCR of (a) *Tc-mlpt*, (b) *Tc-epox* and (c) *Tc-jhamt* in embryos collected 0–6, 6–12, 12–24, 24–48 and 48–72 h after egg laying. Expression was normalized to *Tc-rps3* and is shown relative to the 0–6 h interval. Points represent individual samples (n = 4–6 per interval); horizontal bars show geometric means. Note the log₂ scale

### Embryonic morphology after parental RNAi

Embryos from water-injected females showed normal blastoderm, gastrulation and germ-band stages (Fig. 3a–c). Embryos from *Tc-epox*-injected females included specimens with abnormal anterior curvature and altered head morphology during germ-band elongation; in some of them the presumptive mandibular region was malformed (Fig. 3d–f). Parental RNAi against *Tc-mlpt* produced embryos with additional leg-like appendages on abdominal segments (Fig. 3g, h), as previously described [59]. Embryos from *Tc-jhamt*-injected females showed curved or distorted germ bands with a defective anterior region (Fig. 3i–l). Embryos from *Tc-EcR*-injected females reached germ-band stages with segmented trunks (Fig. 3m, n).

**Fig. 3.**
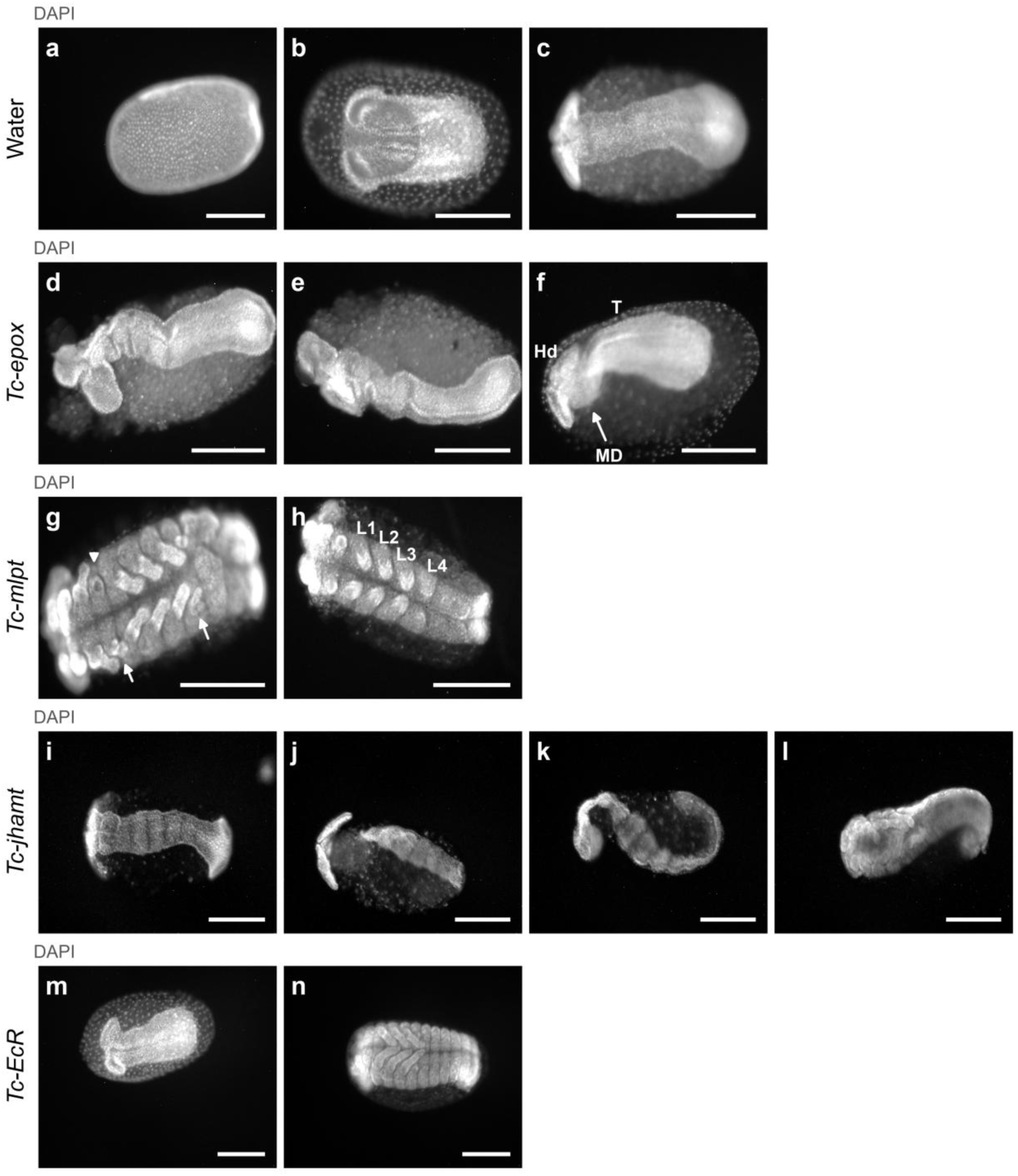
E**m**bryonic **phenotypes after parental RNAi.** (a–c) Embryos from water-injected females at the blastoderm (a), gastrulation (b) and germ-band elongation (c) stages. (d–f) *Tc-epox* RNAi embryos with abnormal anterior curvature and head morphology; the arrow in f indicates the presumptive mandibular region (MD); Hd, head; T, thorax. (g, h) *Tc-mlpt* RNAi embryos with additional leg-like appendages (arrows in g; L1–L4 in h). (i–l) *Tc-jhamt* RNAi embryos with curved or distorted germ bands. (m, n) *Tc-EcR* RNAi embryos at germ-band stages. All panels, DAPI. Scale bars, 200 µm

### *Tc-mlpt* and *Tc-EcR* parental RNAi strongly reduce hatching

Female mortality after injection was 13.3% in both water-control experiments, and 23.3% and 13.3% after *Tc-mlpt* RNAi (mean 18.3%), 26.7% and 40.0% after *Tc-epox* RNAi (mean 33.3%), 30.0% and 53.3% after *Tc-jhamt* RNAi (mean 41.7%) and 26.7% in both experiments after *Tc-EcR* RNAi (Fig. 4a). Hatching of the offspring was strongly reduced after *Tc-mlpt* RNAi (4% and 14% in the two experiments; mean 9.0%) and *Tc-EcR* RNAi (27% and 25%; mean 26.0%), compared with water controls (84% and 64%; mean 74.0%; Fig. 4b). Hatching after *Tc-epox* (84% and 65%; mean 74.5%) and *Tc-jhamt* RNAi (82% and 64%; mean 73.0%) was similar to that of controls in each experiment.

**Fig. 4.**
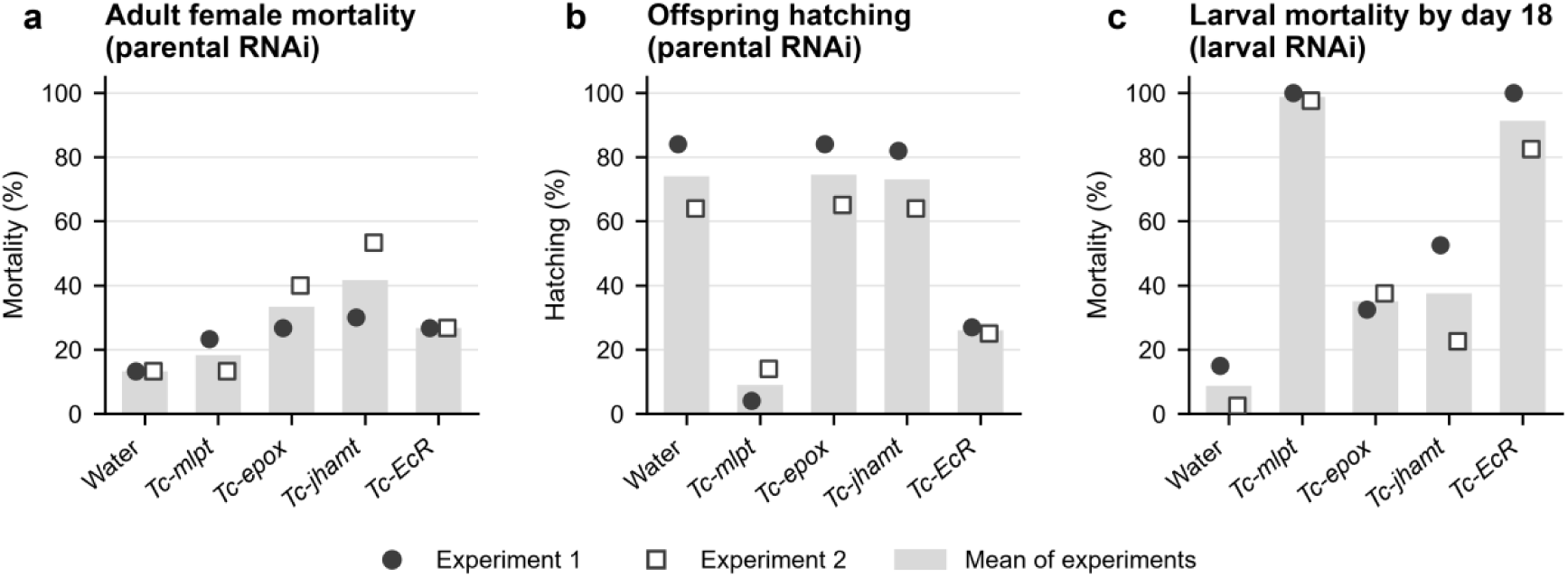
V**i**ability **after parental and larval RNAi.** (a) Mortality of adult females after injection of water or dsRNA. (b) Hatching of eggs laid by injected females (100 eggs per treatment and experiment). (c) Mortality of larvae by day 18 after injection at the third instar (40 larvae per treatment and experiment). Symbols show the two independent experiments; bars show their mean

### Unhatched offspring display treatment-specific cuticle defects

Cuticle preparations of unhatched first-instar larvae revealed distinct defects (Fig. 5); 50 cuticles were examined per treatment. Control cuticles showed a differentiated head capsule, three pairs of thoracic legs and a segmented abdomen (Fig. 5a, a′). As reported above, most eggs from *Tc-epox*- and *Tc-jhamt*-injected females hatched. Fewer than 5% of the *Tc-epox* cuticles were fully segmented but showed abnormal head and posterior terminal structures (Fig. 5b, b′), or had a shortened, swollen body that remained partly enclosed by embryonic membranes (Fig. 5c), and a similar fraction of the *Tc-jhamt* cuticles had compact, rounded bodies with an indistinct head capsule and disorganized appendages (Fig. 5d, e). Although rare, these phenotypes were never observed among the 50 control cuticles.

**Fig. 5.**
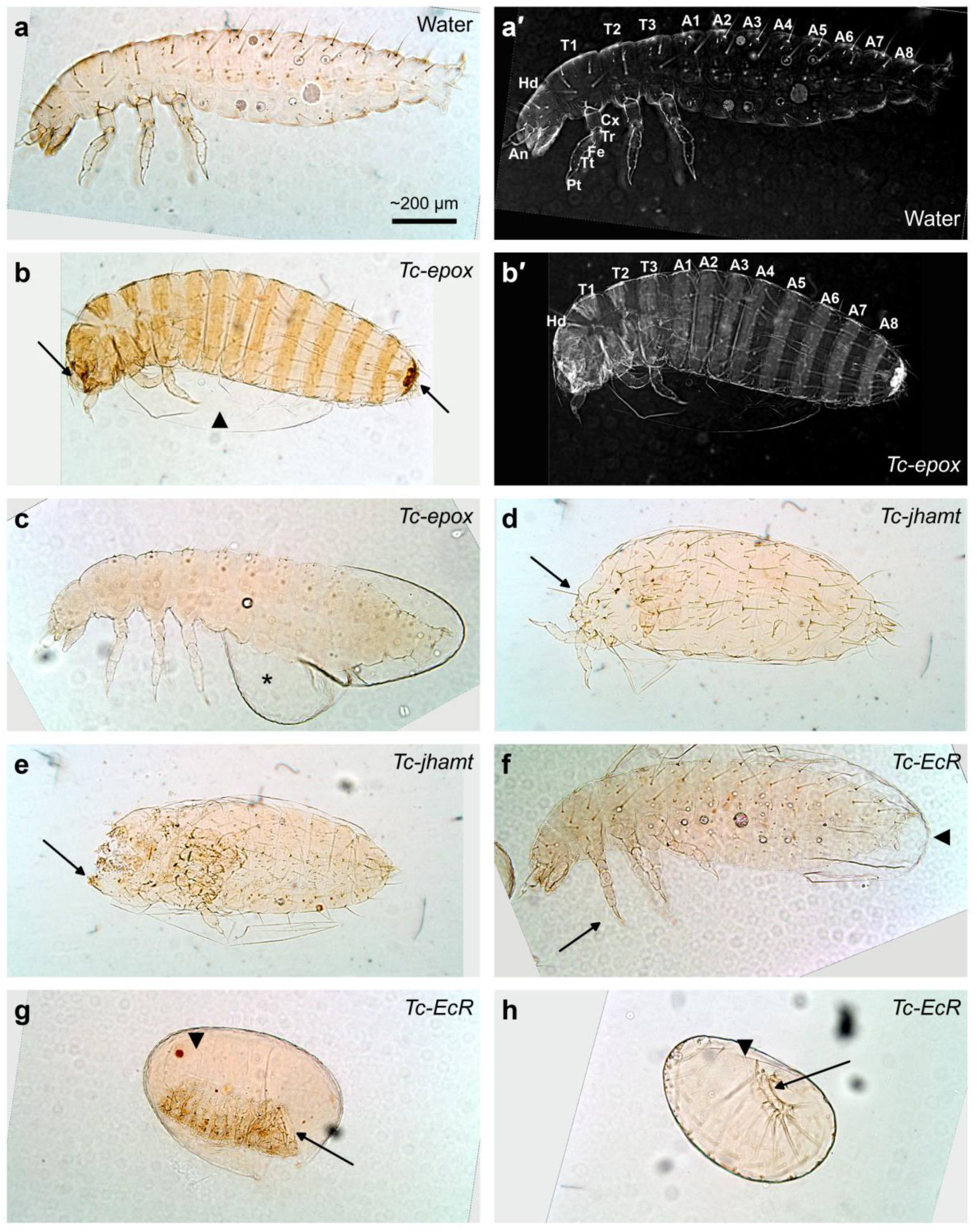
C**u**ticles **of unhatched first-instar larvae after parental RNAi.** Colour bright-field images of cuticle preparations. a′ and b′ show the specimens in a and b as inverted bright-field images, which emphasize setae and segment borders. T1–T3, thoracic segments; A1–A8, abdominal segments; Hd, head; An, antenna; Cx, coxa; Tr, trochanter; Fe, femur; Tt, tibiotarsus; Pt, pretarsus. Arrows indicate the abnormal head and terminal structures (b), the anterior region (d, e), legs (f), the fragmented cuticular band (g) and sparse cuticular elements (h); arrowheads indicate enclosing membranes and the asterisk a membranous sac enclosing the posterior body (c). (a, a′) Control larva with differentiated head, three pairs of thoracic legs and segmented abdomen. (b, b′, c) *Tc-epox*: segmented larvae with abnormal head and terminal structures (b) or a shortened, swollen body partly enclosed by embryonic membranes (c). (d, e) *Tc- jhamt*: compact, rounded larvae with an indistinct head capsule and disorganized appendages. (f–h) *Tc-EcR*: a larva-like cuticle with legs (f), and specimens with an enclosing membrane and a fragmented cuticular band (g) or sparse cuticular elements (h). Larvae are oriented with anterior to the left and dorsal up (a–f); C–F were mirrored to obtain this orientation. All panels are at the same magnification; the bar in a is an approximate 200 µm scale derived as described in the Methods

Among the 50 *Tc-EcR* cuticles examined, we observed a larva-like cuticle with legs (Fig. 5f), a fragmented cuticular band with setae inside an enclosing membrane (Fig. 5g) and sparse cuticular elements (Fig. 5h), indicating severe defects in cuticle formation.

### Ecdysis defects after larval RNAi against *Tc-mlpt* and *Tc-EcR*

Following injection of third-instar larvae, mortality by day 18 was 15.0% and 2.5% in the two water-control experiments (mean 8.8%; Fig. 4c). Mortality reached 100% and 97.5% after *Tc-mlpt* RNAi (mean 98.8%) and 100% and 82.5% after *Tc-EcR* RNAi (mean 91.3%). *Tc-epox* (32.5% and 37.5%; mean 35.0%) and *Tc-jhamt* RNAi (52.5% and 22.5%; mean 37.5%) produced intermediate mortality. Day-by-day counts (Fig. S5) show that deaths after *Tc-mlpt* and *Tc-EcR* RNAi accumulated over the first two weeks, whereas most deaths in the other treatments occurred within the first days after injection, when injection itself is the likely cause.

*Tc-mlpt* larvae died during ecdysis with the old cuticle retained over the anterior or posterior body (Fig. 6c, d, arrows). *Tc-EcR* larvae likewise retained exuviae and showed constrictions of the body (Fig. 6g, h). Larvae that died after *Tc-epox* or *Tc-jhamt* RNAi were darkened and shrunken (Fig. 6e, f).

**Fig. 6.**
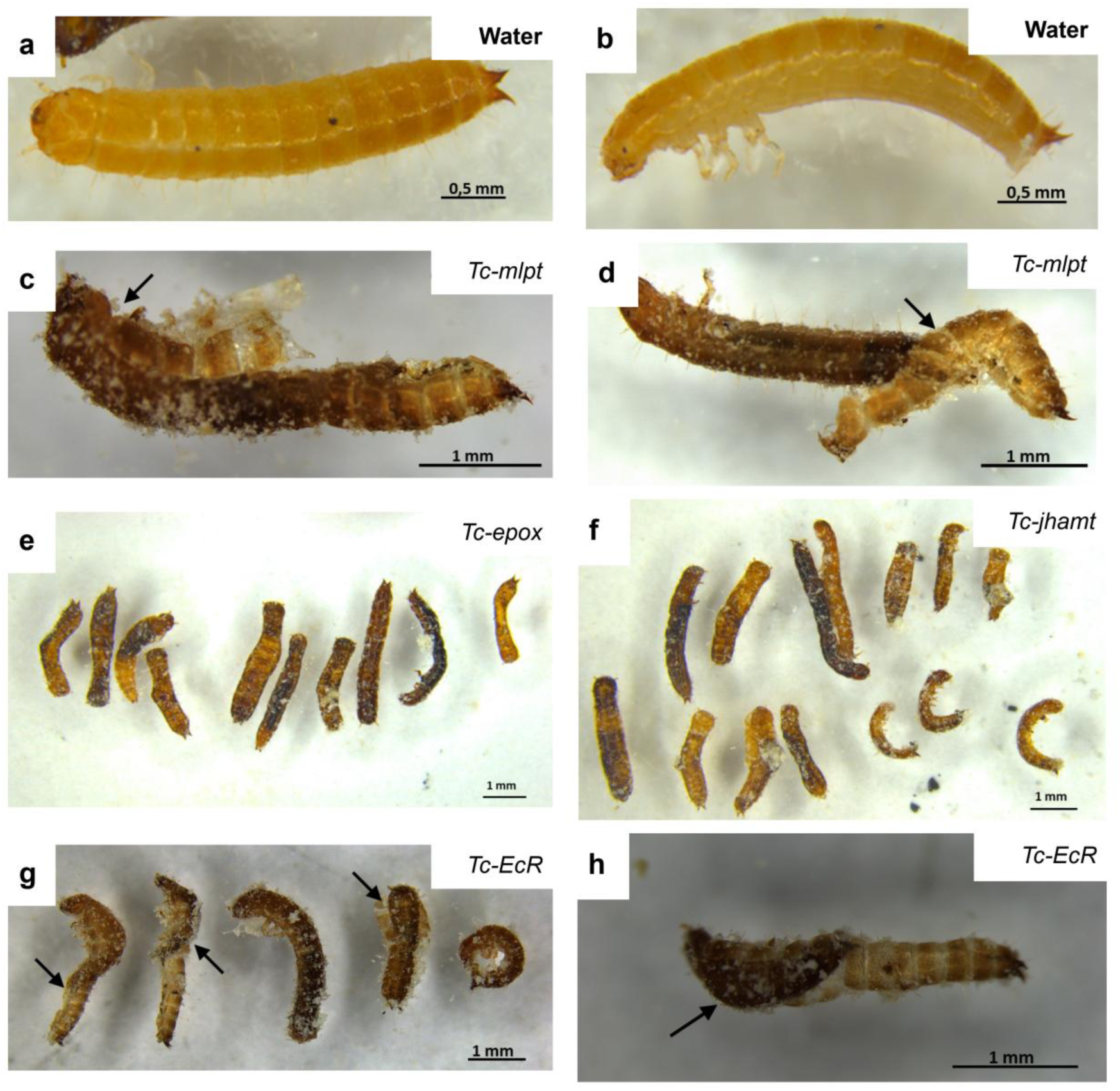
L**a**rval **phenotypes after RNAi.** Larvae injected at the third instar with water (a, b) or dsRNA against *Tc-mlpt* (c, d), *Tc-epox* (e), *Tc-jhamt* (f) or *Tc-EcR* (g, h). *Tc-mlpt* and *Tc-EcR* larvae fail to shed the old cuticle, which remains attached to the anterior or posterior body (arrows). Larvae that died after *Tc-epox* or *Tc-jhamt* RNAi are darkened and shrunken. Scale bars: 0.5 mm (a, b); 1 mm (c–h)

## Discussion

This study shows that genes encoding the final steps of JH biosynthesis are expressed in spatially restricted domains of the *T. castaneum* embryo well before their transcript levels rise at the end of embryogenesis. Parental knockdown of these genes produced morphologically abnormal embryos and cuticles, although mean hatching remained close to that of controls. In contrast, dsRNAs against *Tc-mlpt* and *Tc-EcR* strongly reduced hatching and larval survival, and affected larvae failed to shed the old cuticle.

The late rise of *Tc-jhamt* and *Tc-epox* expression at 48–72 h parallels the late-embryonic JH peak described in the ametabolous firebrat, where JH promotes the switch from morphogenesis to terminal differentiation and is required for hatching [21, 22]. A comparable late embryonic onset of JH action occurs in hemimetabolous embryos, coinciding with deposition of the first nymphal cuticle [23, 24]. Together with the embryonic requirement for Met in *Tribolium* [28], our data are therefore consistent with conservation of a late embryonic JH program across ametabolous, hemimetabolous and at least some holometabolous insects, in contrast to the largely JH-independent embryogenesis reported in *Bombyx* [19]. Measuring JH and methyl farnesoate titers in staged *Tribolium* embryos will be necessary to determine whether the transcriptional rise is accompanied by hormone production, as in *Thermobia*. The rise also coincides with the major embryonic ecdysteroid peak of *Tribolium*. That peak occurs at dorsal closure, at about 120 h of a 153-h embryogenesis at 25°C [43, 44], roughly three quarters of embryonic development. Expressed as fractions of embryogenesis, the weak peak at the extended germ band and the major peak at dorsal closure fall at about 26% and 78% of development. In our cultures at 30°C, where embryogenesis is correspondingly shorter, these fractions correspond approximately to our 12–24 h interval, in which Tc-mlpt expression was maximal, and to our 48–72 h interval. Because the two studies used different temperatures and staging, this correspondence is approximate, and titers and transcripts should be measured in the same staged collections to establish whether the late ecdysteroid peak and the rise of JH biosynthetic gene expression are coordinated.

The early, spatially restricted expression of the biosynthetic genes is more unexpected. In larvae and adults, JHAMT and CYP15 are expressed predominantly in the corpora allata [18, 31, 32], which are not yet differentiated at the blastoderm and germ-band stages examined here, and in *Daphnia* embryos *Jhamt* remains silent until shortly before hatching [35]. The discrete *Tc-epox* domain of the differentiated blastoderm, the bilateral domains of the early germ rudiment and the paired *Tc-jhamt* domains of three trunk segments therefore suggest either local production of sesquiterpenoids in the early embryo or functions of these enzymes independent of systemic JH. In *Drosophila*, which lacks CYP15, *jhamt* and the epoxidase gene *Cyp6g2* have been characterized only in the larval corpora allata [32, 69], and their embryonic expression has not been described. The *Drosophila* corpora allata themselves arise in the embryo from lateral ectodermal cells of the maxillary and labial segments that are serially homologous to the tracheal placodes of the trunk [70]. The segmentally reiterated, lateral position of the *Tc-jhamt* domains is reminiscent of this metameric series, and double labeling with markers of the tracheal and endocrine primordia, such as *trachealess* or *ventral veins lacking*, could test whether early *Tc-jhamt* and *Tc-epox* expression marks precursors of the corpora allata or related segmental cell populations. Transcript localization does not demonstrate colocalization of the two enzymes, substrate availability or hormone synthesis, and double *in situ* hybridization combined with stage-resolved hormone measurements will be needed to test these alternatives.

Parental RNAi against *Tc-jhamt* or *Tc-epox* did not markedly reduce hatching, whereas knockout of *JHAMT* or *CYP15A1* is embryonic lethal in *Thermobia* [22] and parental RNAi against *Met* blocks *Tribolium* embryogenesis [28]. Several non-exclusive explanations may account for this difference. First, parental RNAi may deplete maternal and early zygotic transcripts less efficiently than germline knockout, especially for genes whose expression rises late in embryogenesis. Second, maternally provided JH or precursors may partially compensate for reduced embryonic synthesis. Third, the pathway has built-in redundancy: in *Drosophila*, methyl farnesoate sustains development in the absence of *jhamt* [34], and in *Aedes aegypti* methyl farnesoate supports substantial developmental activity without epoxidation [36]. In *Tribolium*, the duplications that produced *Tc-mt1*, *Tc-mt2* and *Tc-jhamt*, which occurred within the Tenebrionidae (Fig. S1a, d), could add further redundancy if MT1 or MT2 methylate substrates not tested biochemically [33, 37]. The anterior and germ-band defects and malformed cuticles are nevertheless consistent with a contribution of the biosynthetic enzymes to normal embryonic morphogenesis.

These observations bear on the evolution of metamorphosis. Current models propose that JH acted first in embryogenesis and was later co-opted for post-embryonic regulation, first to maintain the nymphal form and, in holometabolous insects, to maintain the larva and time pupal commitment [3, 9, 21]. The pronymph hypothesis further postulates that the holometabolous larva derives from an embryonic stage whose identity is set by JH [1, 5], and molecular changes such as the loss of embryonic *br-C* expression and the reduction of embryonic *E93* [25, 26] are consistent with a re-patterning of the embryonic endocrine program. The conservation of late embryonic biosynthetic-gene expression in *Tribolium* (Fig. 7c) suggests that the late JH program itself was not lost in holometabolous embryos, whereas the early localized domains described here raise the possibility of additional, earlier actions. Whether these act through Met, Kr-h1 and E93, and whether they contribute to specifying larval rather than nymphal features, can be tested by examining *Kr-h1* and *E93* expression after parental knockdown of the biosynthetic genes.

**Fig. 7.**
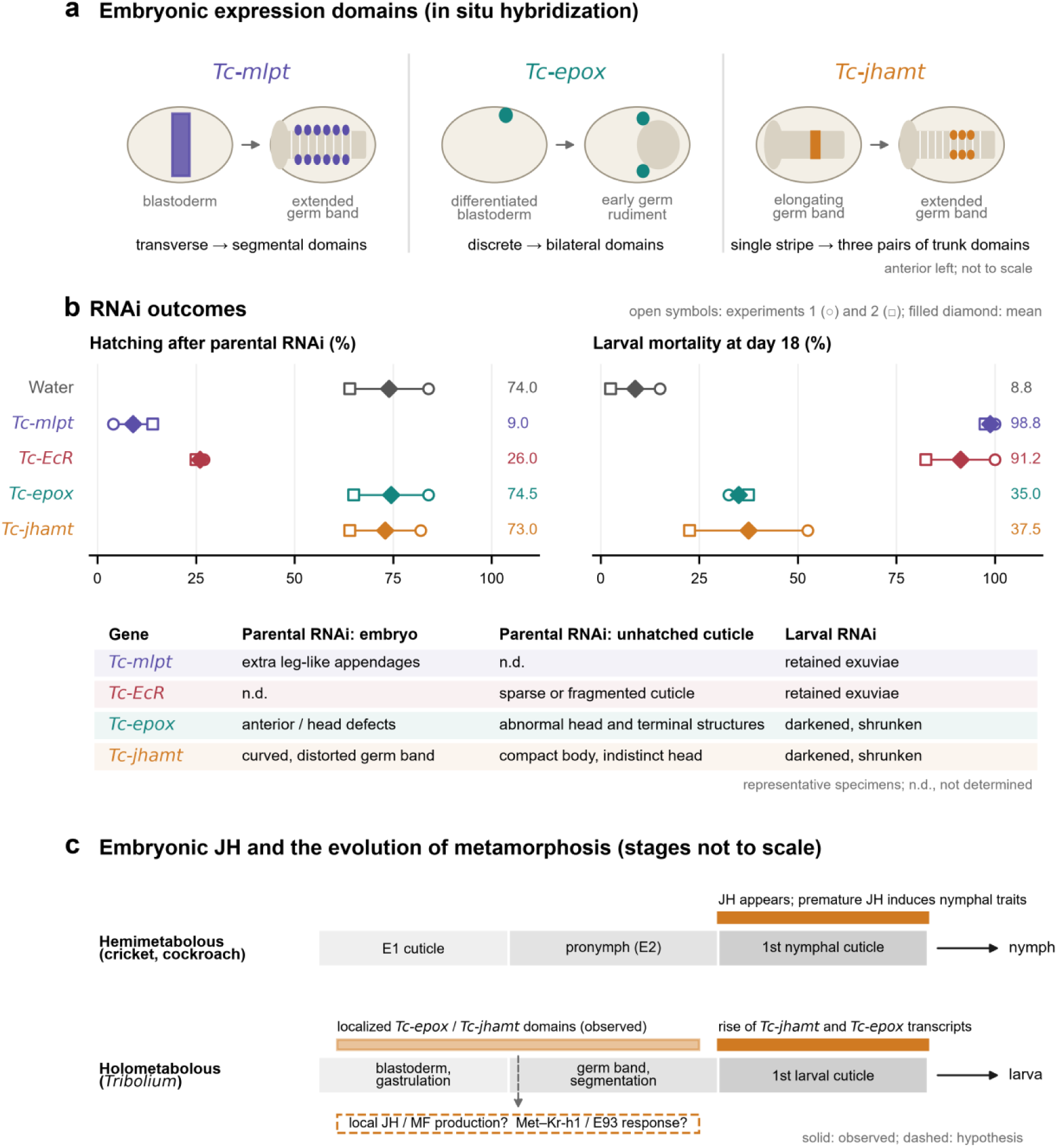
S**u**mmary **of embryonic expression, RNAi outcomes and the timing of embryonic JH.** (a) Schematics of selected *in situ* expression domains of *Tc-mlpt*, *Tc-epox* and *Tc-jhamt* (Fig. 1; Fig. S4); stages are gene-specific examples and are not drawn to scale. (b) Hatching after parental RNAi and mortality by day 18 after larval RNAi (Fig. 4). Open circles and squares show experiments 1 and 2; filled diamonds and numbers show their arithmetic mean. The table summarizes representative morphology (Figs 3, 5 and 6). (c) Timing of embryonic JH relative to embryonic cuticles in hemimetabolous insects (Erezyilmaz et al., 2004; Truman and Riddiford, 2002) compared with the expression of *Tc-epox* and *Tc-jhamt* in *Tribolium* (this study). Solid bars indicate observations; the dashed box and dashed arrows indicate hypotheses that remain to be tested. Stages are not drawn to scale

The cuticle preparations after *Tc-EcR* RNAi revealed severe defects among unhatched offspring, and larvae injected with *Tc-EcR* dsRNA failed to shed the old cuticle. These phenotypes are consistent with the roles of ecdysteroid signaling in embryonic morphogenesis and molting in *Drosophila* and in hemimetabolous insects, where EcR-A knockdown causes molting defects in *Blattella* nymphs [39–41], and with the known requirement of EcR isoforms in *Tribolium* larvae [42]. Because the major embryonic ecdysteroid peak of *Tribolium* coincides with dorsal closure and precedes deposition of the first larval cuticle [44], the sparse or fragmented cuticles of *Tc-EcR* offspring may reflect arrest at or before this stage. Developmental delay or earlier arrest could also produce sparse cuticle, and the *Tc-EcR* dsRNA shares a short sequence with *E75* and *HR38*; an independent, non-overlapping dsRNA will be needed to attribute the embryonic phenotype specifically to EcR.

The larval *Tc-mlpt* phenotype extends the known roles of this gene beyond embryonic segmentation and leg patterning [59, 60]. Linz and Tomoyasu [64] found that *mlpt* RNAi in the last larval instar is lethal just before pupation; our injections at the third instar show that *Tc-mlpt* is already required at the larval–larval molts: affected larvae died during ecdysis with retained exuviae. This phenotype closely resembles the failure of nymphal molting after *mlpt* knockdown in *Rhodnius* [62]. *mlpt* thus appears to have combined an embryonic patterning function [59–61] with a post-embryonic role at each molt since at least the last common ancestor of Paraneoptera and Holometabola. Because the hemipteran-specific smHemiptera peptide has no counterpart in beetles [45, 61], this shared molting function is more likely to depend on the conserved LDPTG-containing peptides, although additional, lineage-specific peptides may have diversified *mlpt* function in hemipterans. In *Drosophila*, Pri peptides mediate ecdysone-dependent temporal programs and epidermal differentiation through Svb processing [49, 51, 53] and, during larval life, repress pupal cuticle genes to set up the soft larval cuticle [57], providing a plausible mechanism for the requirement of *mlpt* in cuticle renewal and ecdysis. Notably, *mlpt* knockdown in *Rhodnius* increased *JHAMT*-like, *EPOX*-like and *Kr-h1* expression [62], suggesting that *mlpt* also influences, directly or indirectly, the expression of JH biosynthetic genes. *Tc-mlpt* and the JH biosynthetic genes showed temporally distinct embryonic profiles in our data, and the maternal expression of *mlpt* in *Aedes* [63] illustrates how its early deployment varies among insects. Measuring *Tc-jhamt*, *Tc-epox* and *Kr-h1* transcript levels after larval *Tc-mlpt* RNAi, and *Tc-mlpt* expression after *Tc-EcR* knockdown, would determine whether *mlpt* integrates ecdysone and JH signaling during beetle molting.

Some limitations should be considered. Parental RNAi affects the injected female as well as her offspring, and increased female mortality after some treatments indicates that maternal effects may contribute to the offspring phenotypes. Quantification of target depletion, a second non-overlapping dsRNA fragment for each gene and phenotype counts from independent cohorts will strengthen the attribution of each phenotype.

In summary, our results provide a spatial and temporal description of JH biosynthetic gene expression in a holometabolous embryo, document severe cuticular defects after *Tc-EcR* RNAi and identify a role for *Tc-mlpt* in larval ecdysis that is shared with the hemimetabolous *Rhodnius* (Fig. 7).

## Materials and methods

### Insect culture and embryo collection

*Tribolium castaneum* was reared on wheat flour at 30°C. Flour was frozen for 24 h at −20°C and heated for 24 h at 50°C before use. For staged collections, adults were allowed to lay eggs in fresh flour, and embryos were collected at 0–6, 6–12, 12–24, 24–48 and 48–72 h after egg laying.

### Phylogenetic analyses

Protein sets were obtained from RefSeq annotations of 25 arthropod genomes (7 Coleoptera, 7 other Holometabola, 5 Hemimetabola, 3 Crustacea and 3 Chelicerata; Table S1). Homologs of Tc-JHAMT (NP_001120783.1) and Tc-CYP15A1 (NP_001308597.1) were identified with DIAMOND v2.2.8 (blastp, --more-sensitive, E < 1e-10), and orthology was assigned by reciprocal best hit against the *T. castaneum* proteome. Partial sequences and proteins shorter than 70% of the query were removed, and one isoform per gene was retained. EcR orthologs were retrieved from RefSeq by annotation and checked by global alignment to Tc-EcR-A. Outgroups were methyltransferases of other families (JHAMT), CYP303a1 and CYP305a1 (CYP15) and human LXRα, LXRβ and FXR (EcR); the JHAMT tree was midpoint-rooted. For a denser analysis of the JHAMT family, the same procedure was applied to 23 RefSeq-annotated beetle genomes (the 7 above and 16 further species, including *Tribolium madens*, *Gnatocerus cornutus* and *Zophobas morio*) and six other insects, after removal of identical or contained sequences; all tenebrionid paralogs and up to eight paralogs per species otherwise were retained, and the tree was midpoint-rooted (Fig. S1d; accession numbers in Supplementary Data). Sequences were aligned with MAFFT v7.526 (L-INS-i), trimmed with trimAl v1.5 (-automated1), and maximum-likelihood trees were inferred with IQ-TREE v2.4.0 using the best-fit model selected by ModelFinder (BIC), with 1,000 ultrafast bootstrap and 1,000 SH-aLRT replicates.

### Probe and dsRNA specificity

Primer pairs (Table S3) were mapped onto their target mRNAs, and the predicted amplicons were compared with all RefSeq transcripts of the *T. castaneum* icTriCast1.1 assembly (GCF_031307605.1). Exact 21-nt matches to other genes were recorded as potential siRNA off-targets, and the best local alignment to any non-target transcript as a measure of potential probe cross-hybridization.

### RNA extraction and RT-PCR

Total RNA was extracted from 100 mg of dechorionated embryos with TRIzol and treated with TURBO DNase. RNA concentration and purity were assessed with a NanoDrop 2000c spectrophotometer, and cDNA was synthesized from 2 µg of total RNA with the High-Capacity cDNA Reverse Transcription Kit (Applied Biosystems). Gene-specific primers were designed with Primer3 (Table S3), and PCR products were analyzed by agarose gel electrophoresis. *Tc-sog*, which is expressed throughout embryogenesis, and *Tc-mlpt* were amplified in the same reactions as positive controls.

### Whole-mount *in situ* hybridization

Embryos were dechorionated in 2% sodium hypochlorite for 3 min and fixed for 30 min in a 1:1 mixture of 4% paraformaldehyde in PBS and heptane. The fixative was replaced with methanol to remove the vitelline membrane, and embryos were stored in methanol at −20°C. After rehydration, embryos were prehybridized for 1 h and hybridized overnight at 55°C with digoxigenin-labeled antisense riboprobes. Signal was detected with alkaline phosphatase-conjugated anti-digoxigenin antibody (1:2000), and nuclei were counterstained with DAPI. Images were acquired on a Leica M205 FA stereomicroscope. Embryos were staged by DAPI morphology (blastoderm, posterior pit, head-lobe/early germ band, elongating and extended germ band) [65] and scored for the presence and position of localized signal. Segment width was measured from the segmental furrows in DAPI images resampled to 1 µm per pixel using the microscope calibration.

### Quantitative RT-PCR

RT-qPCR was performed with SYBR Green chemistry on a StepOnePlus Real-Time PCR System (Applied Biosystems) using primers amplifying 86–125 bp fragments (Table S3). Expression was normalized to *Tc-rps3* [72] with a primer pair whose reverse primer spans an exon–exon junction (125-bp product; Table S3), and expression relative to a calibrator sample from the 0–6 h interval was calculated according to Pfaffl [71]. Four to six samples were analyzed per interval. Values are shown on a log₂ scale with geometric means.

### dsRNA synthesis

Templates for dsRNA synthesis were amplified with gene-specific primers carrying universal adapters, followed by a second PCR with T7-containing universal primers.

dsRNA was transcribed with the T7 RiboMAX Express Large Scale RNA Production System (Promega), precipitated, washed with 70% ethanol and resuspended in injection-grade water at 1 µg/µL.

### Parental RNAi, hatching assays and cuticle preparations

Pupae were sexed and adults were kept separated for three weeks before injection. Thirty females per treatment were immobilized on ice for 10 min and injected laterally into the abdomen with dsRNA or water using a Nanoject II injector (Drummond Scientific), with the largest volume each individual accepted without leakage or rupture; the same criterion was used for the larval injections. After 24 h of recovery at 30°C, males were added at a ratio of one male to four females, and female mortality was recorded. Embryos were collected for fixation, or 100 eggs per treatment were transferred to flour to score hatching; mortality and hatching are reported as percentages of the 30 females and the 100 eggs of each treatment and experiment. Unhatched offspring were dechorionated, mounted in Hoyer’s medium and cleared overnight at 70°C [73], and cuticles were imaged by bright-field microscopy; 50 cuticle preparations were examined per treatment. Each assay was performed in two independent experiments.

### Larval RNAi

Forty third-instar larvae per treatment were injected with dsRNA or water and monitored for 18 days, and dead larvae were counted at each check (Fig. S5; Supplementary Data). Mortality is reported as the number of larvae found dead by day 18 as a percentage of the 40 injected. Two independent experiments were performed. Larvae were photographed on a stereomicroscope to document retained exuviae, body shape and pigmentation.

### Data analysis and figure preparation

RNAi results are reported as the percentage obtained in each of the two experiments and their mean. No statistical tests were applied to these percentages, because each treatment was assayed in two independent experiments. Plots were generated in Python (matplotlib) directly from the source data, which are provided together with the figure scripts (Supplementary Data).

## Acknowledgments

We thank the members of the Laboratório Integrado de Ciências Morfofuncionais and NUPEM/UFRJ for technical assistance and discussions.

## Author contributions

A.S.A. performed experiments, analyzed data and wrote the first draft. L.R., B.R. and D.S. contributed to experimental design, larval RNAi and molecular analyses. R.N.F. conceived and supervised the project, analyzed data and wrote the manuscript. All authors revised and approved the final version.

## Data availability

Source data for all plots (plotted_values.csv, larval_mortality_timecourse.csv), the original Prism files and the scripts used to generate the figures are provided as Supplementary Data.

---

**Fig. S1.** Maximum-likelihood phylogenies of (a) JHAMT-related methyltransferases (midpoint-rooted), (b) CYP15 epoxidases (rooted with CYP303a1 and CYP305a1), (c) ecdysone receptors (rooted with human NR1H receptors) and (d) JHAMT-related methyltransferases from 23 beetle and six other insect genomes (midpoint-rooted; tags give the reciprocal best hit in *T. castaneum* when it is Tc-MT1 or Tc-MT2). Node labels show SH-aLRT/UFBoot support (shown when UFBoot ≥ 50). *T. castaneum* sequences are in bold; colours indicate insect orders and other arthropod groups. Alignments and tree files are provided as Supplementary Data

**Fig. S2.** Position of the *in situ* probe/dsRNA templates and RT-qPCR amplicons on the target mRNAs, and the region shared by *Tc-EcR* isoforms A and B. The *Tc-EcR* template includes 26–28 nt of the DNA-binding domain identical to *E75* and *HR38*

**Fig. S3.** RT-PCR from embryonic cDNA, with Tc-sog as a positive control (a) and Tc- mlpt as a positive control (b). Lane labels are those of the original gels: Puctaq, molecular-weight ladder (sizes in bp, shown at the left of each ladder lane); SOG, Tc-sog; MLPT, Tc-mlpt; EPOX, Tc-epox; ECRa, Tc-EcR; JHAMT and MT3, Tc-jhamt; MT1, Tc-mt1; MT2, Tc-mt2. Amplicon sizes of the target genes are given in Table S3

**Fig. S4.** Calibrated colour images of Tc-epox (a–d) and Tc-jhamt (e–h) expression; primed labels are the corresponding DAPI images. Images were resampled to 1 µm per pixel using the microscope calibration, rotated to align the anteroposterior axis and cropped to a common field. (a) Discrete Tc-epox domain at ∼60% egg length in the differentiated blastoderm; (b) posterior-pit stage; (c) paired bilateral domains in the forming germ rudiment; (d) extended germ band. (e) Tc-jhamt in the blastoderm; (f) early germ band; (g) single transverse trunk stripe during germ-band elongation; (h) paired lateral domains in three consecutive mid-trunk segments, spaced one segment width (55–60 µm) apart. Anterior is to the left. Scale bars, 100 µm

**Fig. S5.** Cumulative number of dead larvae after injection of third-instar larvae with water or dsRNA, in experiment 1 (a) and experiment 2 (b). The dashed line marks day 18, the endpoint used in Fig. 4c. Counts are provided as Supplementary Data

**Table S1.** Accession numbers of all sequences used in the phylogenetic analyses.

**Table S2.** Specificity of probe/dsRNA templates and qPCR amplicons: target transcripts, 21-nt matches to other genes and best alignment to non-target transcripts.

**Table S3.** Primers used for RT-PCR, RT-qPCR and dsRNA template synthesis. Lowercase sequences are universal adapters.

