## Supplementary material for "Embryonic expression and developmental RNAi phenotypes of genes associated with juvenile hormone and ecdysone pathways in *Tribolium castaneum*": FigS1A

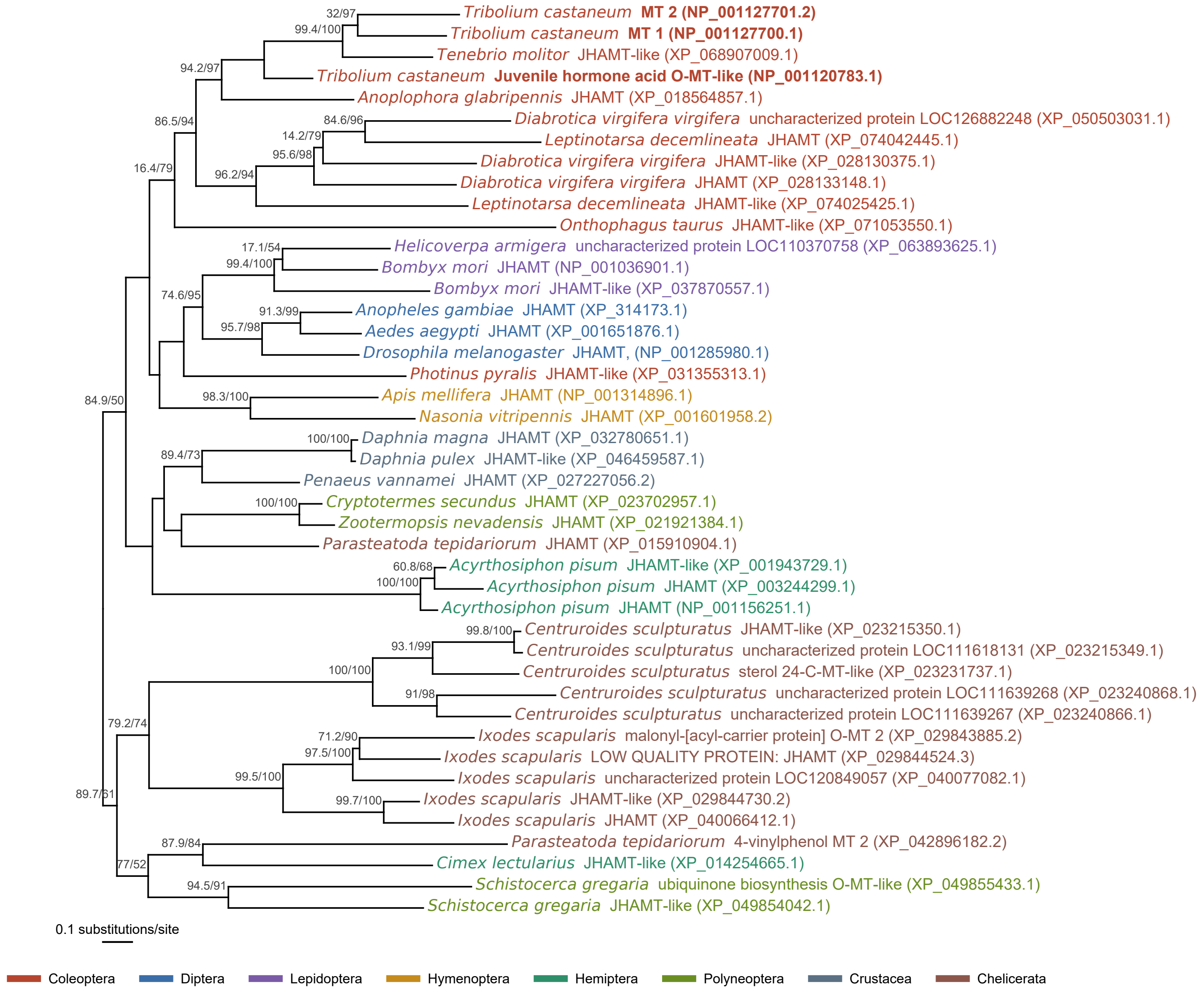
Tribolium castaneum MT 2 (NP_001127701.2)
Tribolium castaneum MT 1 (NP_001127700.1)
Tenebrio molitor JHAMT-like (XP_068907009.1)
Tribolium castaneum Juvenile hormone acid O-MT-like (NP_001120783.1)
Anoplophora glabripennis JHAMT (XP_018564857.1)
Diabrotica virgifera virgifera uncharacterized protein LOC126882248 (XP_050503031.1)
Leptinotarsa decemlineata JHAMT (XP_074042445.1)
Diabrotica virgifera virgifera JHAMT-like (XP_028130375.1)
Diabrotica virgifera virgifera JHAMT (XP_028133148.1)
Leptinotarsa decemlineata JHAMT-like (XP_074025425.1)
Onthophagus taurus JHAMT-like (XP_071053550.1)
Helicoverpa armigera uncharacterized protein LOC110370758 (XP_063893625.1)
Bombyx mori JHAMT (NP_001036901.1)
Bombyx mori JHAMT-like (XP_037870557.1)
Anopheles gambiae JHAMT (XP_314173.1)
Aedes aegypti JHAMT (XP_001651876.1)
Drosophila melanogaster JHAMT, (NP_001285980.1)
Photinus pyralis JHAMT-like (XP_031355313.1)
Apis mellifera JHAMT (NP_001314896.1)
Nasonia vitripennis JHAMT (XP_001601958.2)
Daphnia magna JHAMT (XP_032780651.1)
Daphnia pulex JHAMT-like (XP_046459587.1)
Penaeus vannamei JHAMT (XP_027227056.2)
Cryptotermes secundus JHAMT (XP_023702957.1)
Zootermopsis nevadensis JHAMT (XP_021921384.1)
Parasteatoda tepidariorum JHAMT (XP_015910904.1)
Acyrthosiphon pisum JHAMT-like (XP_001943729.1)
Acyrthosiphon pisum JHAMT (XP_003244299.1)
Acyrthosiphon pisum JHAMT (NP_001156251.1)
Centruroides sculpturatus JHAMT-like (XP_023215350.1)
Centruroides sculpturatus uncharacterized protein LOC111618131 (XP_023215349.1)
Centruroides sculpturatus sterol 24-C-MT-like (XP_023231737.1)
Centruroides sculpturatus uncharacterized protein LOC111639268 (XP_023240868.1)
Centruroides sculpturatus uncharacterized protein LOC111639267 (XP_023240866.1)
Ixodes scapularis malonyl-[acyl-carrier protein] O-MT 2 (XP_029843885.2)
Ixodes scapularis LOW QUALITY PROTEIN: JHAMT (XP_029844524.3)
Ixodes scapularis uncharacterized protein LOC120849057 (XP_040077082.1)
Ixodes scapularis JHAMT-like (XP_029844730.2)
Ixodes scapularis JHAMT (XP_040066412.1)
Parasteatoda tepidariorum 4-vinylphenol MT 2 (XP_042896182.2)
Cimex lectularius JHAMT-like (XP_014254665.1)
Schistocerca gregaria ubiquinone biosynthesis O-MT-like (XP_049855433.1)
Schistocerca gregaria JHAMT-like (XP_049854042.1)
32/97
99.4/100
94.2/97
86.5/94
84.6/96
14.2/79
95.6/98
16.4/79
96.2/94
17.1/54
99.4/100
74.6/95
91.3/99
95.7/98
84.9/50
98.3/100
100/100
89.4/73
100/100
60.8/68
100/100
99.8/100
93.1/99
100/100
91/98
79.2/74
71.2/90
97.5/100
99.5/100
89.7/61
99.7/100
87.9/84
77/52
94.5/91
0.1 substitutions/site
Coleoptera
Diptera
Lepidoptera
Hymenoptera
Hemiptera
Polyneoptera
Crustacea
Chelicerata
