## Supplementary material for "Embryonic expression and developmental RNAi phenotypes of genes associated with juvenile hormone and ecdysone pathways in *Tribolium castaneum*": FigS1B

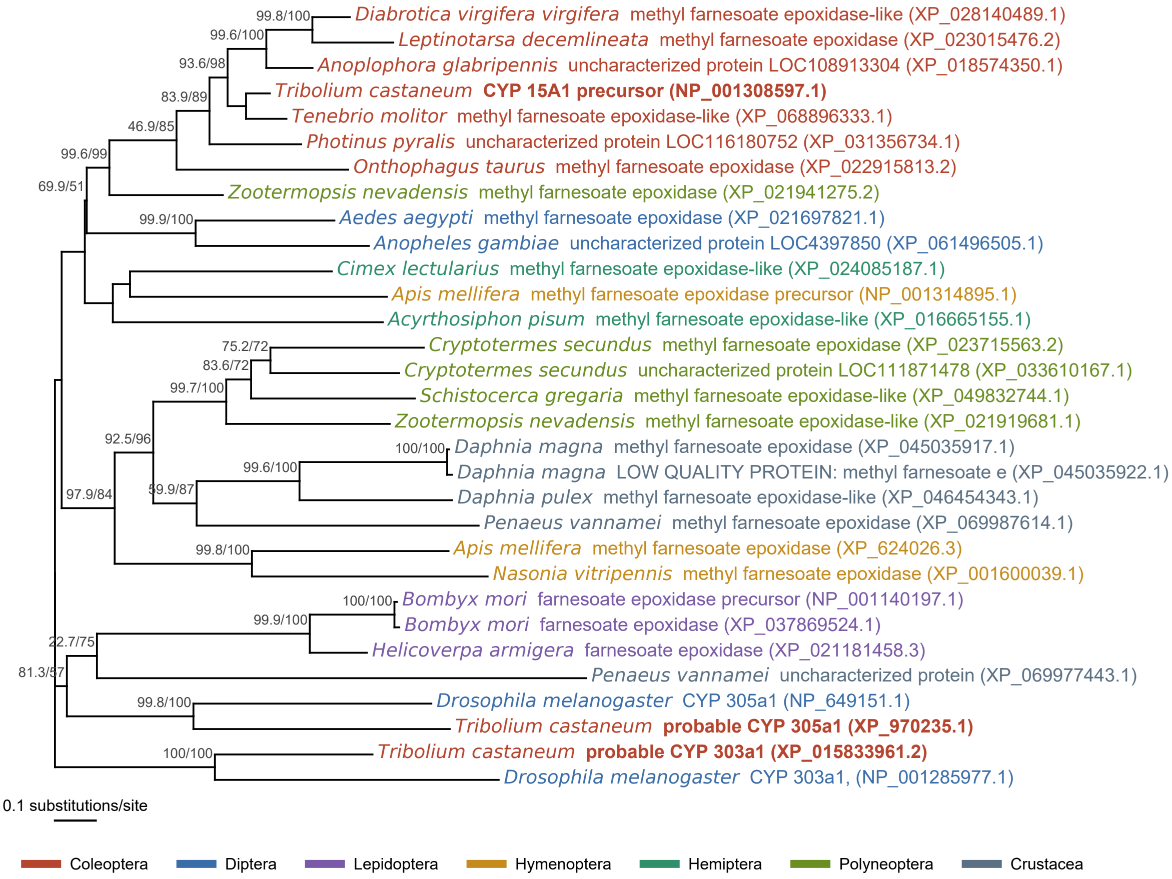

99.8/100
Diabrotica virgifera virgifera methyl farnesoate epoxidase-like (XP_028140489.1)
99.6/100
Leptinotarsa decemlineata methyl farnesoate epoxidase (XP_023015476.2)
93.6/98
Anoplophora glabripennis uncharacterized protein LOC108913304 (XP_018574350.1)
Tribolium castaneum CYP 15A1 precursor (NP_001308597.1)
83.9/89
Tenebrio molitor methyl farnesoate epoxidase-like (XP_068896333.1)
46.9/85
Photinus pyralis uncharacterized protein LOC116180752 (XP_031356734.1)
99.6/99
Onthophagus taurus methyl farnesoate epoxidase (XP_022915813.2)
69.9/51
Zootermopsis nevadensis methyl farnesoate epoxidase (XP_021941275.2)
99.9/100
Aedes aegypti methyl farnesoate epoxidase (XP_021697821.1)
Anopheles gambiae uncharacterized protein LOC4397850 (XP_061496505.1)
Cimex lectularius methyl farnesoate epoxidase-like (XP_024085187.1)
Apis mellifera methyl farnesoate epoxidase precursor (NP_001314895.1)
Acyrthosiphon pisum methyl farnesoate epoxidase-like (XP_016665155.1)
75.2/72
Cryptotermes secundus methyl farnesoate epoxidase (XP_023715563.2)
83.6/72
Cryptotermes secundus uncharacterized protein LOC111871478 (XP_033610167.1)
99.7/100
Schistocerca gregaria methyl farnesoate epoxidase-like (XP_049832744.1)
Zootermopsis nevadensis methyl farnesoate epoxidase-like (XP_021919681.1)
92.5/96
100/100
Daphnia magna methyl farnesoate epoxidase (XP_045035917.1)
99.6/100
Daphnia magna LOW QUALITY PROTEIN: methyl farnesoate e (XP_045035922.1)
97.9/84
59.9/87
Daphnia pulex methyl farnesoate epoxidase-like (XP_046454343.1)
Penaeus vannamei methyl farnesoate epoxidase (XP_069987614.1)
99.8/100
Apis mellifera methyl farnesoate epoxidase (XP_624026.3)
Nasonia vitripennis methyl farnesoate epoxidase (XP_001600039.1)
100/100
Bombyx mori farnesoate epoxidase precursor (NP_001140197.1)
99.9/100
Bombyx mori farnesoate epoxidase (XP_037869524.1)
22.7/75
Helicoverpa armigera farnesoate epoxidase (XP_021181458.3)
81.3/57
Penaeus vannamei uncharacterized protein (XP_069977443.1)
99.8/100
Drosophila melanogaster CYP 305a1 (NP_649151.1)
Tribolium castaneum probable CYP 305a1 (XP_970235.1)
100/100
Tribolium castaneum probable CYP 303a1 (XP_015833961.2)
Drosophila melanogaster CYP 303a1, (NP_001285977.1)
0.1 substitutions/site
Coleoptera
Diptera
Lepidoptera
Hymenoptera
Hemiptera
Polyneoptera
Crustacea
