## Supplementary material for "Embryonic expression and developmental RNAi phenotypes of genes associated with juvenile hormone and ecdysone pathways in *Tribolium castaneum*": FigS1C

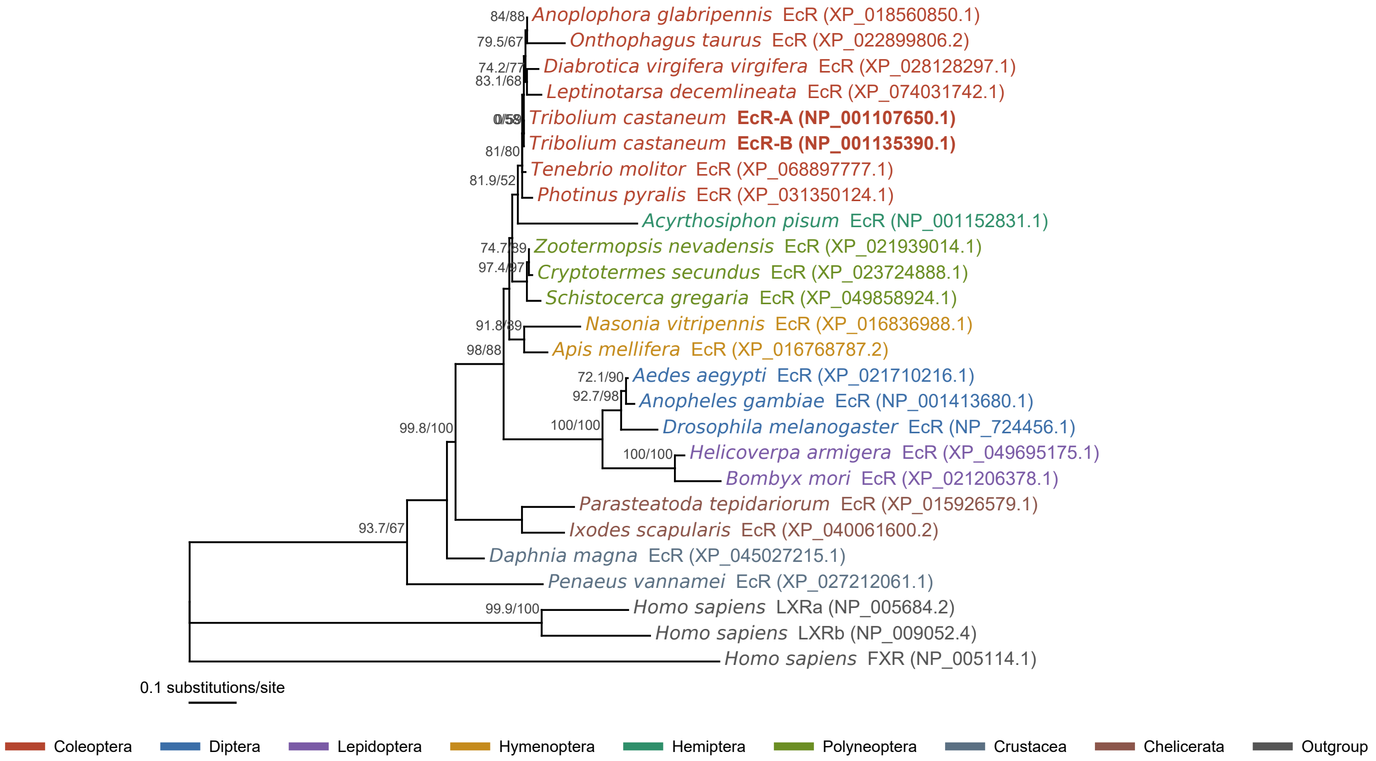

84/88
Anoplophora glabripennis EcR (XP_018560850.1)
79.5/67
Onthophagus taurus EcR (XP_022899806.2)
74.2/77
Diabrotica virgifera virgifera EcR (XP_028128297.1)
83.1/68
Leptinotarsa decemlineata EcR (XP_074031742.1)
0/58
Tribolium castaneum EcR-A (NP_001107650.1)
Tribolium castaneum EcR-B (NP_001135390.1)
81/80
Tenebrio molitor EcR (XP_068897777.1)
81.9/52
Photinus pyralis EcR (XP_031350124.1)
Acyrthosiphon pisum EcR (NP_001152831.1)
74.7/89
Zootermopsis nevadensis EcR (XP_021939014.1)
97.4/97
Cryptotermes secundus EcR (XP_023724888.1)
Schistocerca gregaria EcR (XP_049858924.1)
91.8/89
Nasonia vitripennis EcR (XP_016836988.1)
98/88
Apis mellifera EcR (XP_016768787.2)
72.1/90
Aedes aegypti EcR (XP_021710216.1)
92.7/98
Anopheles gambiae EcR (NP_001413680.1)
100/100
Drosophila melanogaster EcR (NP_724456.1)
99.8/100
100/100
Helicoverpa armigera EcR (XP_049695175.1)
Bombyx mori EcR (XP_021206378.1)
Parasteatoda tepidariorum EcR (XP_015926579.1)
93.7/67
Ixodes scapularis EcR (XP_040061600.2)
Daphnia magna EcR (XP_045027215.1)
Penaeus vannamei EcR (XP_027212061.1)
99.9/100
Homo sapiens LXRa (NP_005684.2)
Homo sapiens LXRb (NP_009052.4)
Homo sapiens FXR (NP_005114.1)
0.1 substitutions/site
Coleoptera
Diptera
Lepidoptera
Hymenoptera
Hemiptera
Polyneoptera
Crustacea
Chelicerata
Outgroup
