## Supplementary material for "Embryonic expression and developmental RNAi phenotypes of genes associated with juvenile hormone and ecdysone pathways in *Tribolium castaneum*": FigS1D

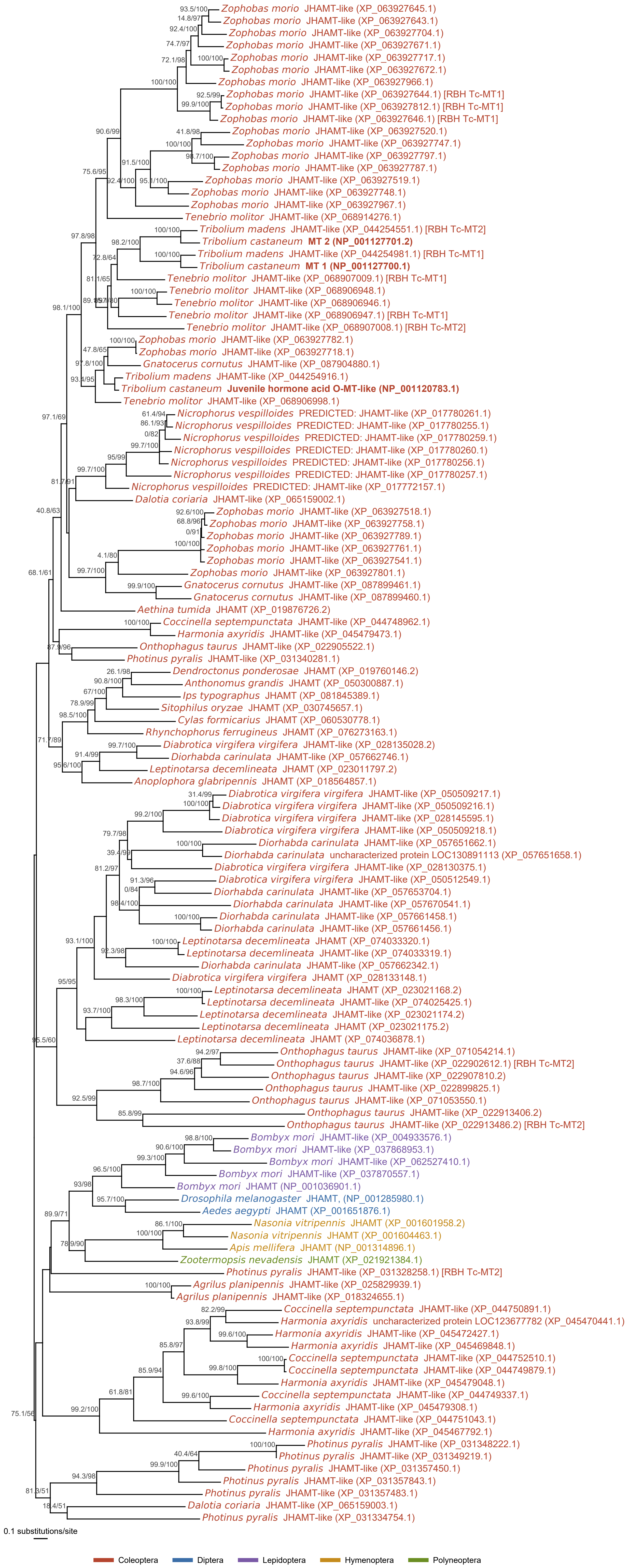
Zophobas morio JHAMT-like (XP_063927645.1)
Zophobas morio JHAMT-like (XP_063927643.1)
Zophobas morio JHAMT-like (XP_063927704.1)
Zophobas morio JHAMT-like (XP_063927671.1)
Zophobas morio JHAMT-like (XP_063927717.1)
Zophobas morio JHAMT-like (XP_063927672.1)
Zophobas morio JHAMT-like (XP_063927966.1)
Zophobas morio JHAMT-like (XP_063927644.1) [RBH Tc-MT1]
Zophobas morio JHAMT-like (XP_063927812.1) [RBH Tc-MT1]
Zophobas morio JHAMT-like (XP_063927646.1) [RBH Tc-MT1]
Zophobas morio JHAMT-like (XP_063927520.1)
Zophobas morio JHAMT-like (XP_063927747.1)
Zophobas morio JHAMT-like (XP_063927797.1)
Zophobas morio JHAMT-like (XP_063927787.1)
Zophobas morio JHAMT-like (XP_063927519.1)
Zophobas morio JHAMT-like (XP_063927748.1)
Zophobas morio JHAMT-like (XP_063927967.1)
Tenebrio molitor JHAMT-like (XP_068914276.1)
Tribolium madens JHAMT-like (XP_044254551.1) [RBH Tc-MT2]
Tribolium castaneum MT 2 (NP_001127701.2)
Tribolium madens JHAMT-like (XP_044254981.1) [RBH Tc-MT1]
Tribolium castaneum MT 1 (NP_001127700.1)
Tenebrio molitor JHAMT-like (XP_068907009.1) [RBH Tc-MT1]
Tenebrio molitor JHAMT-like (XP_068906948.1)
Tenebrio molitor JHAMT-like (XP_068906946.1)
Tenebrio molitor JHAMT-like (XP_068906947.1) [RBH Tc-MT1]
Tenebrio molitor JHAMT-like (XP_068907008.1) [RBH Tc-MT2]
Zophobas morio JHAMT-like (XP_063927782.1)
Zophobas morio JHAMT-like (XP_063927718.1)
Gnatocerus cornutus JHAMT-like (XP_087904880.1)
Tribolium madens JHAMT-like (XP_044254916.1)
Tribolium castaneum Juvenile hormone acid O-MT-like (NP_001120783.1)
Tenebrio molitor JHAMT-like (XP_068906998.1)
Nicrophorus vespilloides PREDICTED: JHAMT-like (XP_017780261.1)
Nicrophorus vespilloides PREDICTED: JHAMT-like (XP_017780255.1)
Nicrophorus vespilloides PREDICTED: JHAMT-like (XP_017780259.1)
Nicrophorus vespilloides PREDICTED: JHAMT-like (XP_017780260.1)
Nicrophorus vespilloides PREDICTED: JHAMT-like (XP_017780256.1)
Nicrophorus vespilloides PREDICTED: JHAMT-like (XP_017780257.1)
Nicrophorus vespilloides PREDICTED: JHAMT-like (XP_017772157.1)
Dalotia coriaria JHAMT-like (XP_065159002.1)
Zophobas morio JHAMT-like (XP_063927518.1)
Zophobas morio JHAMT-like (XP_063927758.1)
Zophobas morio JHAMT-like (XP_063927789.1)
Zophobas morio JHAMT-like (XP_063927761.1)
Zophobas morio JHAMT-like (XP_063927541.1)
Zophobas morio JHAMT-like (XP_063927801.1)
Gnatocerus cornutus JHAMT-like (XP_087899461.1)
Gnatocerus cornutus JHAMT-like (XP_087899460.1)
Aethina tumida JHAMT (XP_019876726.2)
Coccinella septempunctata JHAMT-like (XP_044748962.1)
Harmonia axyridis JHAMT-like (XP_045479473.1)
Onthophagus taurus JHAMT-like (XP_022905522.1)
Photinus pyralis JHAMT-like (XP_031340281.1)
Dendroctonus ponderosae JHAMT (XP_019760146.2)
Anthonomus grandis JHAMT (XP_050300887.1)
Ips typographus JHAMT (XP_081845389.1)
Sitophilus oryzae JHAMT (XP_030745657.1)
Cylas formicarius JHAMT (XP_060530778.1)
Rhynchophorus ferrugineus JHAMT (XP_076273163.1)
Diabrotica virgifera virgifera JHAMT-like (XP_028135028.2)
Diorhabda carinulata JHAMT-like (XP_057662746.1)
Leptinotarsa decemlineata JHAMT (XP_023011797.2)
Anoplophora glabripennis JHAMT (XP_018564857.1)
Diabrotica virgifera virgifera JHAMT-like (XP_050509217.1)
Diabrotica virgifera virgifera JHAMT-like (XP_050509216.1)
Diabrotica virgifera virgifera JHAMT-like (XP_028145595.1)
Diabrotica virgifera virgifera JHAMT-like (XP_050509218.1)
Diorhabda carinulata JHAMT-like (XP_057651662.1)
Diorhabda carinulata uncharacterized protein LOC130891113 (XP_057651658.1)
Diabrotica virgifera virgifera JHAMT-like (XP_028130375.1)
Diabrotica virgifera virgifera JHAMT-like (XP_050512549.1)
Diorhabda carinulata JHAMT-like (XP_057653704.1)
Diorhabda carinulata JHAMT-like (XP_057670541.1)
Diorhabda carinulata JHAMT-like (XP_057661458.1)
Diorhabda carinulata JHAMT-like (XP_057661456.1)
Leptinotarsa decemlineata JHAMT (XP_074033320.1)
Leptinotarsa decemlineata JHAMT-like (XP_074033319.1)
Diorhabda carinulata JHAMT-like (XP_057662342.1)
Diabrotica virgifera virgifera JHAMT (XP_028133148.1)
Leptinotarsa decemlineata JHAMT (XP_023021168.2)
Leptinotarsa decemlineata JHAMT-like (XP_074025425.1)
Leptinotarsa decemlineata JHAMT-like (XP_023021174.2)
Leptinotarsa decemlineata JHAMT (XP_023021175.2)
Leptinotarsa decemlineata JHAMT (XP_074036878.1)
Onthophagus taurus JHAMT-like (XP_071054214.1)
Onthophagus taurus JHAMT-like (XP_022902612.1) [RBH Tc-MT2]
Onthophagus taurus JHAMT-like (XP_022907810.2)
Onthophagus taurus JHAMT-like (XP_022899825.1)
Onthophagus taurus JHAMT-like (XP_071053550.1)
Onthophagus taurus JHAMT-like (XP_022913406.2)
Onthophagus taurus JHAMT-like (XP_022913486.2) [RBH Tc-MT2]
Bombyx mori JHAMT-like (XP_004933576.1)
Bombyx mori JHAMT-like (XP_037868953.1)
Bombyx mori JHAMT-like (XP_062527410.1)
Bombyx mori JHAMT-like (XP_037870557.1)
Bombyx mori JHAMT (NP_001036901.1)
Drosophila melanogaster JHAMT, (NP_001285980.1)
Aedes aegypti JHAMT (XP_001651876.1)
Nasonia vitripennis JHAMT (XP_001601958.2)
Nasonia vitripennis JHAMT (XP_001604463.1)
Apis mellifera JHAMT (NP_001314896.1)
Zootermopsis nevadensis JHAMT (XP_021921384.1)
Photinus pyralis JHAMT-like (XP_031328258.1) [RBH Tc-MT2]
Agrilus planipennis JHAMT-like (XP_025829939.1)
Agrilus planipennis JHAMT-like (XP_018324655.1)
Coccinella septempunctata JHAMT-like (XP_044750891.1)
Harmonia axyridis uncharacterized protein LOC123677782 (XP_045470441.1)
Harmonia axyridis JHAMT-like (XP_045472427.1)
Harmonia axyridis JHAMT-like (XP_045469848.1)
Coccinella septempunctata JHAMT-like (XP_044752510.1)
Coccinella septempunctata JHAMT-like (XP_044749879.1)
Harmonia axyridis JHAMT-like (XP_045479048.1)
Coccinella septempunctata JHAMT-like (XP_044749337.1)
Harmonia axyridis JHAMT-like (XP_045479308.1)
Coccinella septempunctata JHAMT-like (XP_044751043.1)
Harmonia axyridis JHAMT-like (XP_045467792.1)
Photinus pyralis JHAMT-like (XP_031348222.1)
Photinus pyralis JHAMT-like (XP_031349219.1)
Photinus pyralis JHAMT-like (XP_031357450.1)
Photinus pyralis JHAMT-like (XP_031357843.1)
Photinus pyralis JHAMT-like (XP_031357483.1)
Dalotia coriaria JHAMT-like (XP_065159003.1)
Photinus pyralis JHAMT-like (XP_031334754.1)
0.1 substitutions/site
Coleoptera
Diptera
Lepidoptera
Hymenoptera
Polyneoptera
