## Supplementary material for "Embryonic expression and developmental RNAi phenotypes of genes associated with juvenile hormone and ecdysone pathways in *Tribolium castaneum*": FigS2

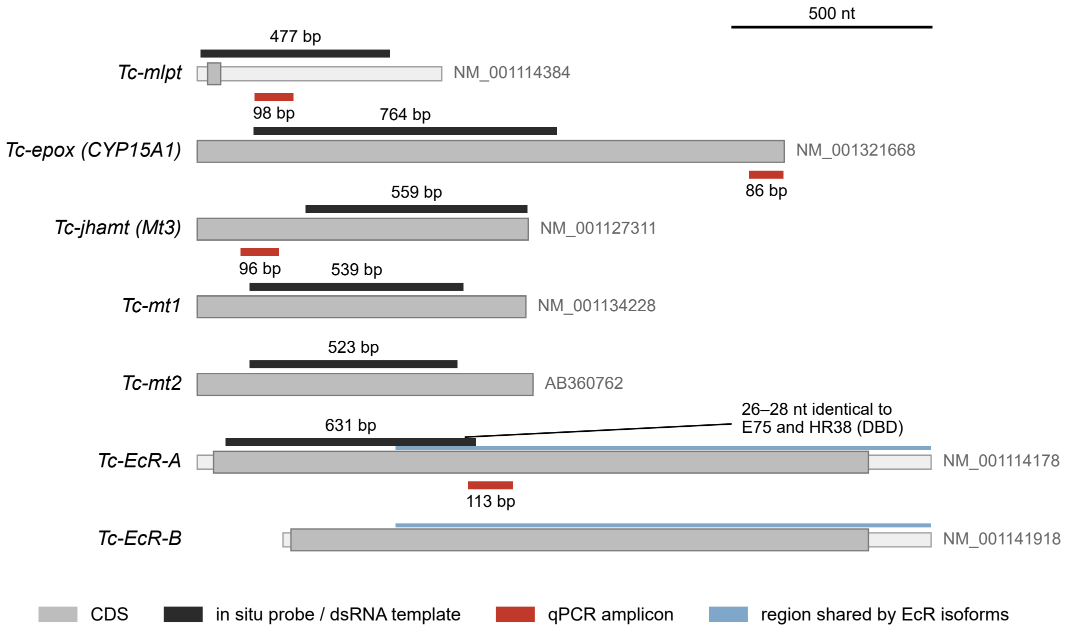

500 nt
477 bp
Tc-mlpt
NM_001114384
98 bp
764 bp
Tc-epox (CYP15A1)
NM_001321668
86 bp
559 bp
Tc-jhamt (Mt3)
NM_001127311
96 bp
539 bp
Tc-mt1
NM_001134228
523 bp
Tc-mt2
AB360762
26–28 nt identical to
E75 and HR38 (DBD)
631 bp
Tc-EcR-A
NM_001114178
113 bp
Tc-EcR-B
NM_001141918
CDS
in situ probe / dsRNA template
qPCR amplicon
region shared by EcR isoforms
