## Supplementary figures and images for "Embryonic expression and developmental RNAi phenotypes of genes associated with juvenile hormone and ecdysone pathways in *Tribolium castaneum*"

### FigS3

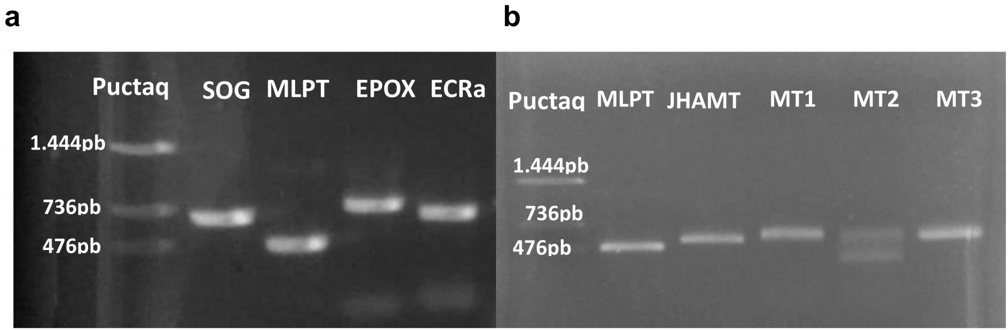
a
b
Puctaq
SOG
MLPT
EPOX
ECRa
1.444pb
736pb
476pb
Puctaq
MLPT
JHAMT
MT1
MT2
MT3
1.444pb
736pb
476pb

### FigS4

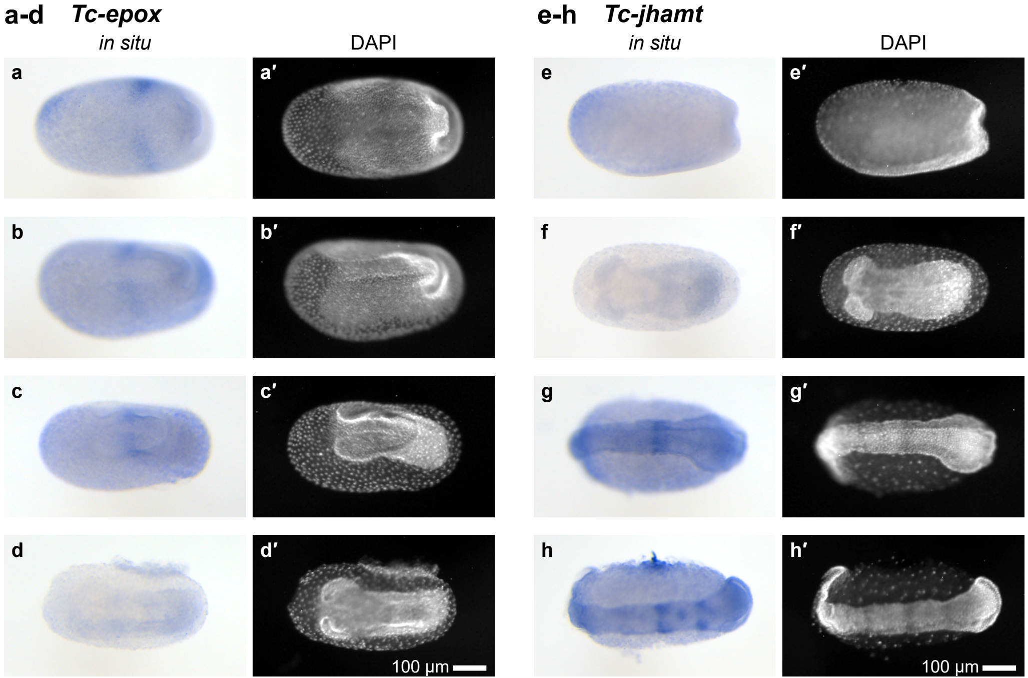
a-d Tc-epox
in situ
DAPI
e-h Tc-jhamt
in situ
DAPI
a
a′
b
b′
c
c′
d
d′
100 µm
e
e′
f
f′
g
g′
h
h′
100 µm
