## Supplementary material for "Embryonic expression and developmental RNAi phenotypes of genes associated with juvenile hormone and ecdysone pathways in *Tribolium castaneum*": FigS5

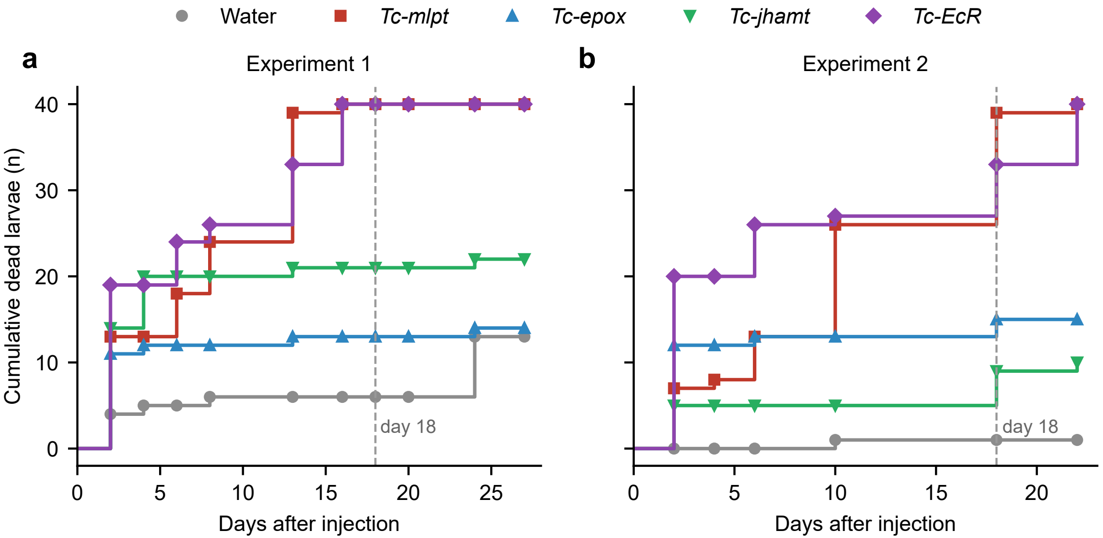

Water
Tc-mlpt
Tc-epox
Tc-jhamt
Tc-EcR
a
Experiment 1
b
Experiment 2
Cumulative dead larvae (n)
40
30
20
10
0
day 18
day 18
0
5
10
15
20
25
0
5
10
15
20
Days after injection
Days after injection
